# Two novel heteropolymer-forming proteins maintain multicellular shape of the cyanobacterium *Anabaena* sp. PCC 7120

**DOI:** 10.1101/553073

**Authors:** Benjamin L. Springstein, Dennis J. Nürnberg, Christian Woehle, Julia Weissenbach, Marius L. Theune, Andreas O. Helbig, Iris Maldener, Tal Dagan, Karina Stucken

**Author notes:** Corresponding authors: BLS, KS. Present address: Department of Microbiology, Blavatnick Institute, Harvard Medical School, Boston, MA, USA. Present address: Institute of Experimental Physics, Free University of Berlin, Berlin, Germany (Dennis J. Nürnberg). Present address: Max Planck-Genome-centre cologne, Max Planck Institute for Plant Breeding Research, Cologne, Germany (Christian Woehle). Present address: Faculty of Biology, Technion-Israel Institute of Technology, Haifa, Israel (Julia Weissenbach).

## Abstract

Polymerizing and filament-forming proteins are instrumental for numerous cellular processes such as cell division and growth. Their function in stabilization and localization of protein complexes and replicons is achieved by a filamentous structure. Known filamentous proteins assemble into homopolymers consisting of single subunits – e.g. MreB and FtsZ in bacteria – or heteropolymers that are composed of two subunits, e.g. keratin and α/β tubulin in eukaryotes. Here, we describe two novel coiled-coil-rich proteins (CCRPs) in the filament forming cyanobacterium *Anabaena* sp. PCC 7120 (hereafter *Anabaena*) that assemble into a heteropolymer and function in the maintenance of the *Anabaena* multicellular shape (termed trichome). The two CCRPs – Alr4504 and Alr4505 (named ZicK and ZacK) – are strictly interdependent for the assembly of protein filaments *in vivo* and polymerize nucleotide-independently *in vitro*, similar to known intermediate filament (IF) proteins. A ΔzicKΔzacK double mutant is characterized by a zigzagged cell arrangement and hence a loss of the typical linear *Anabaena* trichome shape. ZicK and ZacK interact with themselves, with each other, with the elongasome protein MreB, the septal junction protein SepJ and the divisome associate septal protein SepI. Our results suggest that ZicK and ZacK function in cooperation with SepJ and MreB to stabilize the *Anabaena* trichome and are likely essential for the manifestation of the multicellular shape in *Anabaena*. Our study reveals the presence of filament-forming IF-like proteins whose function is achieved through the formation of heteropolymers in cyanobacteria.

## Introduction

Cytoskeletal proteins that polymerize to form protein filaments are paramount in bacterial cell biology where they play a role in cell division, alignment of bacterial microcompartments (BMCs), chromosome and plasmid segregation, organization of cell polarity and the determination of cell shape (Wagstaff and Löwe, 2018). For example, FtsZ (Van De Putte *et al.*, 1964; de Boer *et al.*, 1992), the prokaryotic homolog to the eukaryotic tubulin (Löwe and Amos, 1998; Nogales *et al.*, 1998), is a main component of the divisome (den Blaauwen *et al.*, 2017), a multiprotein complex that governs cell division in bacteria and self-assembles into a proteinaceous ring (called Z-ring) at the midcell position (Bi and Lutkenhaus, 1991). Another key bacterial cytoskeletal protein is MreB (Jones *et al.*, 2001), which is a homolog of the eukaryotic actin (de Boer *et al.*, 1992; Ent *et al.*, 2001) and a crucial component of the multi-protein complexes termed the elongasome. This complex modulates cell elongation in many rod-shaped bacteria through regulating peptidoglycan (PG) biogenesis (Errington and Wu, 2017). Both, FtsZ and MreB monomers assemble into filamentous strands (protofilaments), consisting of only one type of monomer, termed homopolymers (Wagstaff and Löwe, 2018). The cell division in prokaryotes markedly contrasts the division of plastid organelles in photosynthetic eukaryotes that are of cyanobacterial origin (e.g., (Dagan *et al.*, 2013)). Cell division in plastids is dependent on the cooperative function and heteropolymerization of two FtsZ homologs, FtsZ1 and FtsZ2 in the green lineages and FtsZA and FtsZB in the red lineage. However, each FtsZ homolog is also self-sufficient to form homopolymers (reviewed by (Chen *et al.*, 2018)). In contrast, a likely horizontally transferred pair of tubulin homologs, BtubA and BtubB from *Prothescobacter* spp., exclusively assembles into heteropolymers *in vitro* (Schlieper *et al.*, 2005), revealing similar properties than eukaryotic microtubules that are heteropolymers composed of α and β tubulin monomers (Alberts *et al.*, 2014). Eukaryotic IF proteins, despite sharing substantially the same building blocks and a high degree of coiled-coil (CC) domains (Fuchs and Weber, 1994), which are considered excellent mediators of protein-protein interactions (Mason and Arndt, 2004), only form heteropolymers with a subset of other IF proteins within their same assembly group but otherwise form obligate homopolymers (Herrmann and Aebi, 2000).

Polymer-forming coiled-coil-rich proteins (CCRPs) have been shown to play a role also in multicellularity traits in myxobacteria and actinomycetes (reviewed by (Lin and Thanbichler, 2013; Wagstaff and Löwe, 2018)). Similar to eukaryotic IFs (Fuchs and Weber, 1994), many bacterial CCRPs perform cytoskeletal functions through their ability to self-assemble into filaments *in vivo* and *in vitro* in a self-sufficient and co-factor independent manner (Ausmees *et al.*, 2003; Bagchi *et al.*, 2008; Specht *et al.*, 2011; Holmes *et al.*, 2013). The CCRP Crescentin determines the *C. crescentus* typical curved morphology by aligning to the inner cell curvature and exuding local mechanical constrains on the PG biosynthesis, likely through cooperation with MreB (Ausmees *et al.*, 2003; Charbon *et al.*, 2009). In analogy to eukaryotic IF proteins, Crescentin assembles into straight protein filaments with a width of 10 nm and displays a similar domain organization (Ausmees *et al.*, 2003). However, while Crescentin is often considered a prokaryotic homologue to eukaryotic IF proteins, its restricted distribution to only one identified organism questions real homologous relationships and rather suggests that it was acquired by horizontal gene transfer (Erickson, 2007; Wickstead and Gull, 2011). Multicellular actinobacteria, such as *Streptomyces* spp., grow by building new cell wall (*i.e*., PG) only at the cell poles, independent of MreB (Letek *et al.*, 2008), a striking different cell growth than in most other bacteria (Surovtsev and Jacobs-Wagner, 2018). This characteristic polar growth mode is organized by a cytoskeletal network of at least three CCRPs - DivIVA, Scy and FilP – that directly interact with each other to form the polarisome (Holmes *et al.*, 2013). FilP and Scy, independently self-assemble into filaments *in vitro* (Bagchi *et al.*, 2008; Holmes *et al.*, 2013; Javadi *et al.*, 2019), thereby fulfilling major IF-like criteria (Wagstaff and Löwe, 2018). *In vivo*, however, Scy does not form filaments and instead accumulates as foci at future branching points (Holmes *et al.*, 2013), while FilP localizes as gradient-like caudates at the hyphal tips (Fröjd and Flärdh, 2019), instead of forming distinct filaments as observed for Crescentin (Ausmees *et al.*, 2003). Although of essential importance for growth and cell shape, the polarisome does not directly regulate multicellularity in Actinobacteria, which is instead maintained by the highly reproducible and coordinated formation of Z-ring ladders during sporulation (Schwedock *et al.*, 1997; Claessen *et al.*, 2014).

Among prokaryotes, Cyanobacteria exhibit the largest morphological diversity, comprising unicellular species as well as complex cell-differentiating multicellular species (Rippka *et al.*, 1979). For the model multicellular cyanobacterium *Anabaena*, it is imperative to form stable trichomes in order to cope with external influences such as shearing stress (Corrales-Guerrero *et al.*, 2013; Flores *et al.*, 2016). Under nitrogen-deprived growth conditions, *Anabaena* develops specialized cell types for nitrogen fixation (heterocysts), which are evenly spaced among the *Anabaena* trichome and provide other vegetative cells with fixed nitrogen compounds like glutamine (Herrero *et al.*, 2016). In *Anabaena*, proteinaceous cell-joining structures that allow intercellular transport (*i.e*., cell-cell communication) and function by gating are termed septal junctions (analogous to eukaryotic gap junctions; (Wilk *et al.*, 2011; Flores *et al.*, 2018; Weiss *et al.*, 2019)). Septal junctions consist of several structural elements, an intracellular cap, a plug inside the cytoplasmic membrane formed by the septal junction protein FraD and a tube traversing the septum through nanopores in the peptidoglycan (Weiss et al. 2019). The correct positioning of the septal protein SepJ, which is involved in septum maturation and filament stability, among others, dependents on the FtsZ-driven divisome component FtsQ (Ramos-León *et al.*, 2015), which links the early and late assembly components of the divisome (Choi *et al.*, 2018). FtsZ was shown to be essential for *Anabaena* viability and to assemble in a typical Z-ring structure at future septum sites in vegetative cells while being downregulated in heterocysts (Zhang *et al.*, 1995; Sakr *et al.*, 2006a; Klint *et al.*, 2007). In contrast, MreB is dispensable for *Anabaena* viability but determines the typical rectangular-like, since Δ*mreB* mutant cells show pronounced rounded and swollen morphotype. Unlike in many unicellular bacteria, MreB does not affect chromosome segregation, which was found to be governed, at least in part, by random segregation in *Anabaena* (Hu *et al.*, 2007). Maintenance of the rectangular cell shape is furthermore dependent on a class B penicillin-binding-protein (PBP) (Burnat *et al.*, 2014) and AmiC-type cell wall amidases in *Anabaena* (Bornikoel *et al.*, 2017; Kieninger *et al.*, 2019), suggesting that loss of normal cell shape is commonly associated with defects in PG biogenesis (Fenton *et al.*, 2016).

In this work, we aimed to identify proteins that play a role in cyanobacterial morphology and multicellularity. Searching for IF-like CCRPs, we identified two novel CCRPs in *Anabaena* that are capable of assembly exclusively into a heteropolymer *in vitro* and *in vivo* and that have a putative role in the *Anabaena* linear trichome shape.

## Results

### CCRPs ZicK and ZacK from *Anabaena* are conserved in Cyanobacteria

A computational survey of the *Anabaena* genome for protein-coding genes containing a high coiled-coil content (Springstein *et al.*, 2020b) revealed two CCRPs Alr4504 and Alr4505; here we term the two CCRPs ZicK and ZacK, respectively (that is, zig and zag in German). ZicK is predicted to contain five distinct coiled-coil (CC) domains while ZacK has four CC domains (Fig 1A; Supplementary File 1). Using PSORTb (v.3.0.2), both ZicK and ZacK are predicted to be cytoplasmic proteins, which is corroborated by the absence of detectable transmembrane domains (predicted using TMHMM v. 2.0). Since both proteins (and their homologs) are annotated as hypothetical proteins (Supplementary File 2), we validated their transcription under standard (BG11) and diazotrophic (BG11_0_) growth conditions (Supplementary Fig. 1B,C). The genomic neighbourhood of *zicK* and *zacK* motivated us to test for a common transcriptional regulation of both genes (*i.e*., an operon structure), however, we did not identify a common transcript (using RT-PCR; Supplementary Fig. 1A,B). Searching for known proteins sharing structural similarities to ZicK/ZacK using I-TASSER revealed structural similarities between ZicK and the eukaryotic cytolinker protein plectin, and of ZacK with the cell division protein EzrA, a predicted structural similarity that was previously associated with other bacterial CCRPs, including Crescentin, HmpF_Syn_ and HmpF_Syc_ (Springstein *et al.*, 2020b). Further annotation using the NCBI conserved domain search (Marchler-Bauer *et al.*, 2016) showed that ZicK and ZacK contain “structural maintenance of chromosomes” (SMC) domains (Fig. 1B), similarly to what we previously identified in other self-polymerizing cyanobacterial CCRPs (Springstein *et al.*, 2020b). A search for ZicK/ZacK homologs by sequence similarity revealed that they are absent in picocyanobacteria (*i.e*., *Synechococcus*/*Prochlorococcus*) and generally rare in unicellular cyanobacteria. Otherwise, about 50% of the examined 168 cyanobacterial genomes have homologs for the two genes (Fig. 1C and Supplementary File 2). Several heterocystous cyanobacteria lacking ZacK/ZacK homologs are characterized by a reduced genome (e.g., *Nostoc azollae* PCC 0708 and *Richelia intracellularis*). Additionally, *Chlorogloeopsis* spp. that forms multiseriate filaments is lacking the homologs and several strains of *Fischerella* spp., which forms true branching filaments, harbour only a ZacK homolog. The protein sequence of ZicK and ZacK homologs is well conserved, with about 55% amino acids identity among the homologs. The number of CC domains, however, differ among ZicK and ZacK homologs: between 3-6 CC domains in ZicK homologs, and 3-10 domains in ZacK homologs (Supplementary File 1). Notably, ZicK and ZacK are neighbours in 53 out 72 genomes; both proteins and their genomic neighbourhood is highly conserved among heterocystous cyanobacteria (Fig. 1C,D).

**Fig. 1:**
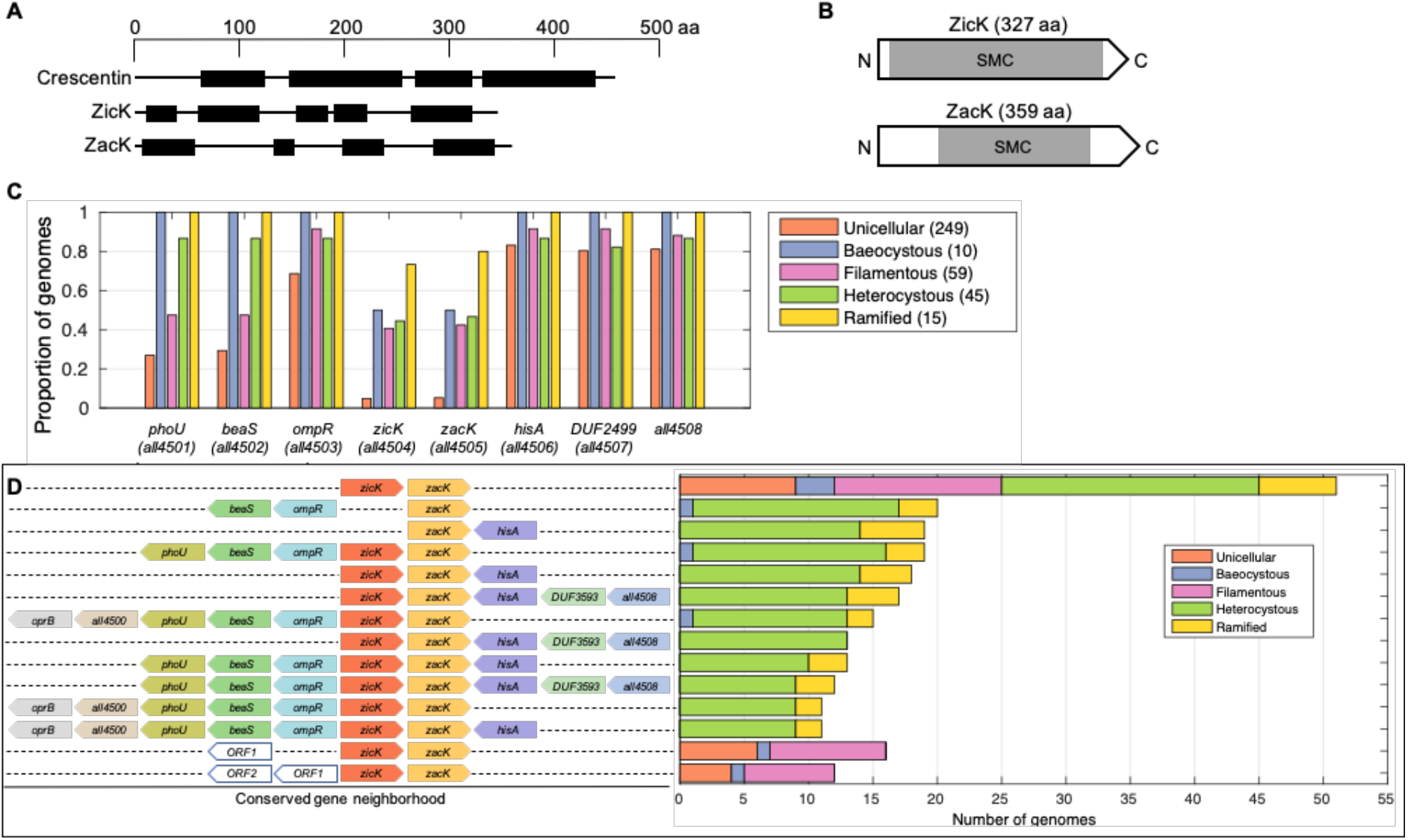
Conservation of ZicK and Zack among cyanobacterial species and domain architecture. (**A**) Depiction of coiled-coil domains of ZicK and ZacK as identified by the COILS algorithm (Lupas *et al.*, 1991) with a window width of 21. As a comparison, Crescentin from *C. crescentus* is also included. The scale on top is given in amino acid residues (aa) and amino acid sequences in coiled-coil conformation are depicted by black bars, while non-coiled-coil sequences are represented by black lines. (**B**) Schematic depiction of the domain architecture of ZicK and ZacK. The SMC domains predicted for both proteins are depicted by grey bars. (**C**) The presence of homologs of ZicK/ZacK in cyanobacteria main types (total number of genomes in the analysis is shown in the legend). Note that the presence of ZicK/ZacK homologs in baeocystous cyanobacteria is in accordance with a recent suggestion of a multicellular ancestry of species in that group (Urrejola *et al.*, 2020). (**D**) Genomic neighbourhood of ZicK/ZacK in cyanobacterial genomes. Conserved syntenic blocks (CSBs; *i.e*., conserved gene order) are shown on the left; the number of genomes where the same gene order has been identified is shown by a bar right of the conserved gene order. Genes with clear annotation or unique locus name are designated as ORF. Note that the CSBs are not mutually exclusive - *i.e*., the longer CSBs where ZicK/ZacK are neighbours (and in the same orientation) include the ZicK/ZacK CSB (1st line on top). The 2nd line from the top shows the CSB in organisms where ZicK is absent. The two CSBs at the bottom reveal that the genomic neighbourhood of ZicK/ZacK in non-heterocystous cyanobacteria is different in comparison to heterocystous cyanobacteria.

### ZicK and ZacK are interdependent for polymerization *in vitro*

As a prerequisite for proteins to be considered as IF-like proteins, it is imperative for them to be able to self-interact *in vivo* and to polymerize into long protein filaments *in vitro* (Wagstaff and Löwe, 2018). To investigate the *in vitro* polymerization properties of ZicK and ZacK, we employed an *in vitro* polymerization assay that we previously established to test CCRPs’ polymerization properties (Springstein *et al.*, 2020b). As a positive control for our approach, we used Crescentin (Fig. 1A), which formed an extensive filamentous network in our *in vitro* assay (Supplementary Fig. 2). As negative controls, we included empty vector-carrying *E. coli* cells, GroEL1.2 from *Chlorogloeopsis fritschii* PCC 6912 (known to form oligomers (Weissenbach *et al.*, 2017)) and the highly soluble maltose binding protein (MBP), all of which were tested negative for filament formation *in vitro* using our approach (Supplementary Fig. 2). Purified ZicK protein formed into amorphous, non-filamentous protein aggregates while ZacK assembled into aggregated sheet-like structures (Fig. 2A). The vast majority of ZacK protein precipitated into clumps of aggregates upon renaturation, which resembled the structures observed for GroEL1.2, suggesting that ZacK has only a partial capacity to self-polymerize or, more likely, is highly unstable on its own *in vitro*. Inspired by the close genomic localization of *zicK* and *zacK,* we next tested for a potential heteropolymerization of both proteins. This revealed that ZicK and ZacK co-assembled into a meshwork of protein heteropolymers upon co-renaturation (Fig. 2A, Supplementary Fig. 3). While both, ZicK and ZacK renatured alone, formed aggregates in the dialysis tubes that were detectable with the naked eye (similar to our observations from GroEL1.2), co-renatured ZicK/ZacK remained in solution, a common property of eukaryotic IFs (Köster *et al.*, 2015). Next, we tested whether the co-filamentation was dosage dependent and observed that distinct protein filaments could be detected *in vitro* only with equal amounts of ZicK and ZacK (Supplementary Fig. 3). To further test for an *in vivo* self-interaction, we analysed the self-binding capacity ZicK and ZacK using the bacterial adenylate cyclase two-hybrid (BACTH) assay and found that ZicK and ZacK interact with themselves and also with each other (Fig. 2B) confirming the heterologous binding capacity of both proteins. Consequently, ZicK and ZacK fulfil a major characteristic of IF and IF-like proteins as they are able to self-assemble into filament-like structures *in vitro*, although, unlike other bacterial IF-like CCRPs, this assembly exclusively occurs as a heteropolymer.

**Fig. 2:**
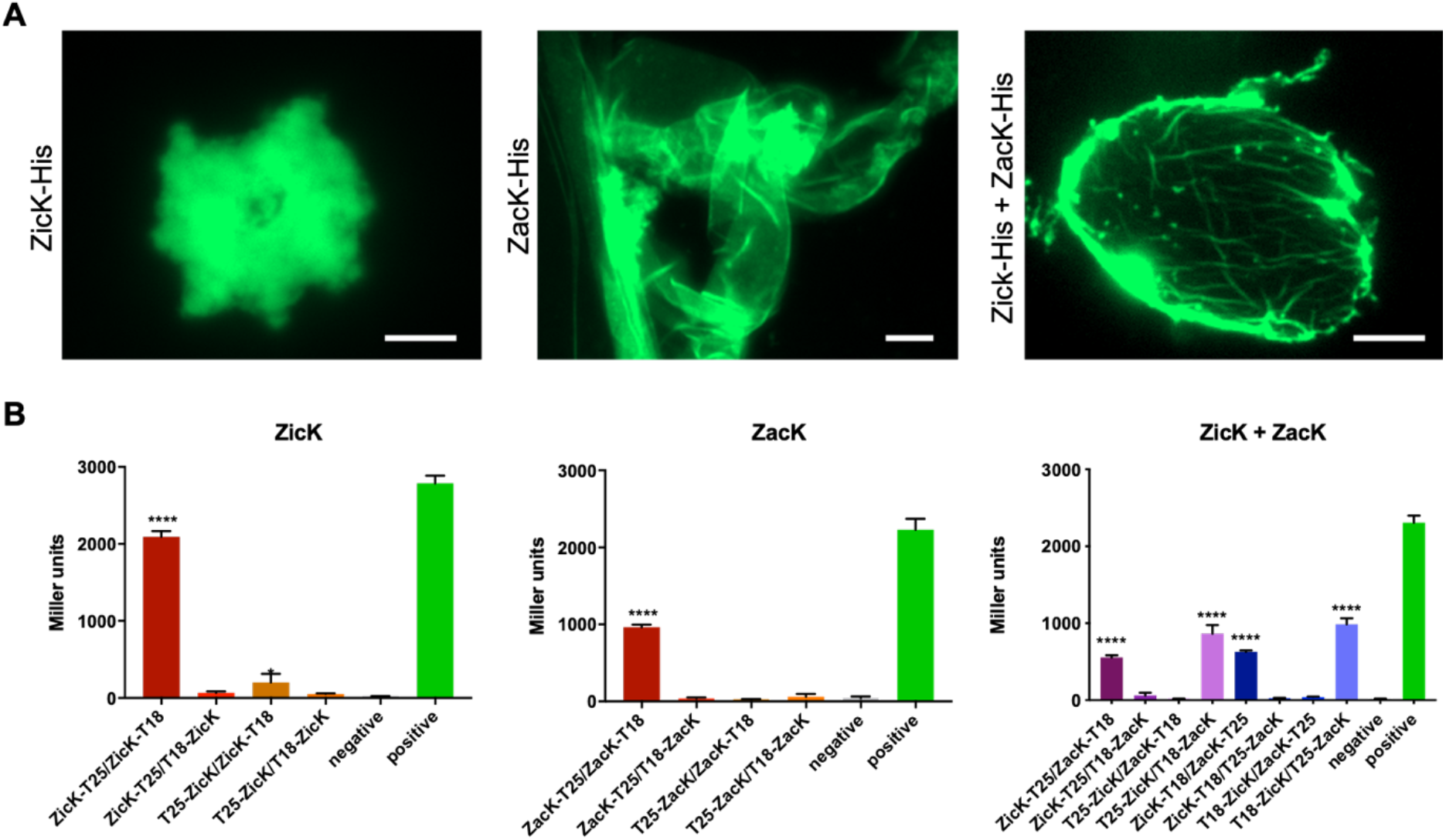
ZicK and ZacK form heteropolymer filament-like structures and interact *in vivo*. (**A**) Epifluorescence micrographs of NHS-Fluorescein-stained *in vitro* structures formed by purified and renatured ZicK-His (1 mg ml^−1^), ZacK-His (0.5 mg ml^−1^) or co-renatured ZicK-His and ZacK-His (0.25 mg ml^−1^ each) in 25 mM HEPES, pH 7.4 (ZacK) or HLB (ZicK and co-renatured ZicK/ZacK) renaturation buffer. Note: although ZacK formed somewhat filamentous structures *in vitro*, the vast majority of ZacK clumped into aggregates, reminiscent of GroEL1.2 (Supplementary Fig 2). (**B**) BACTH assays of *E. coli* cells co-expressing indicated T25 and T18 translational fusions of all possible pair-wise combinations of ZicK and ZacK. *E. coli* cells were subjected to beta-galactosidase assay in triplicates from three independent colonies grown for 2 d at 20°C. Quantitative values are given in Miller units, and the mean results from three independent colonies are presented. Negative: N-terminal T25 fusion construct of the respective protein co-transformed with empty pUT18C. Positive: Zip/Zip control. Error bars indicate standard deviations (n = 3). Values indicated with asterisks are significantly different from the negative control. ****: p < .0001 (Student‘s t-test).

### ZicK and ZacK are interdependent for polymerization *in vivo*

To examine the *in vivo* localization pattern of ZicK and ZacK, we initially expressed translational GFP fusions of both proteins from the replicative pRL25C plasmid, which is commonly used in experimental work in *Anabaena* (Sakr *et al.*, 2006b; Sakr *et al.*, 2006a; Hu *et al.*, 2007; Du *et al.*, 2012). The expression of ZicK-GFP and ZacK-GFP from their respective native promoters (as predicted using BPROM (Solovyev and Salamov, 2011)) revealed no discernible expression of ZacK-GFP, while ZicK-GFP accumulated within the cells as dot-like aggregates (Supplementary Fig. 4A). We assume that the lack of ZacK-GFP expression from its predicted native promoter is based on the uncertainty of the precise promoter site prediction. Alternatively, and although not expressed as an operon, expression of *zacK* could be affected by the expression of *zicK* by so far unknown mechanisms. Consequently, we proceeded to investigate the *in vivo* localization of both proteins from the copper-regulated *petE* promoter (P_petE_), which is commonly used to study the subcellular protein localization in *Anabaena* (e.g., FtsZ, MreB and SepI; (Sakr *et al.*, 2006a; Sakr *et al.*, 2006b; Hu *et al.*, 2007; Springstein *et al.*, 2020a). The expression of ZicK-GFP and ZacK-GFP from P_petE_ in *Anabaena* independently did not reveal distinct or filamentous structures but resulted in the formation of inclusion body-like aggregates within the cells (Fig. 3A), similar to those observed for P_zicK_::*zicK-gfp* (Supplementary Fig. 4A). We could not detect any structures when we expressed ZicK and ZacK N-terminally fused to GFP, suggesting that the N-terminus is key for proper protein folding. Consequently, we next proceeded to co-express the two proteins with different fluorophores (from P_petE_): ZicK C-terminally fused to eCFP (ZicK-eCFP) and ZacK C-terminally fused to GFP (ZacK-GFP). This revealed the formation of a ZicK/ZacK heteropolymer filament-like structure that usually localized along the longitudinal cell axis and in rare occasions also perpendicular to the cell axis (Fig. 3B). The prominent formation of the ZicK/ZacK heteropolymer was furthermore evident as electron-dense filament-like structures in ultrathin sections using electron microscopy (Fig. 3C). To confirm that the localization of fluorophore tagged ZicK and ZacK is not affected by the wild type (WT) *zicK* or *zacK* alleles, we additionally localized both proteins individually or together in a Δ*zicK*Δ*zacK* double mutant. These experiments revealed the same localization pattern as in *Anabaena* WT, suggesting that natively present ZicK or ZacK proteins do not affect the formation of ZicK-GFP, ZacK-GFP or the ZicK-eCFP/ZacK-GFP heteropolymer (Supplementary Fig. 4B,C). The intracellular localization of the ZicK/ZacK heteropolymer in *Anabaena* indicates that the polymer is either anchored at the cell poles or specifically split during cell division, as ZicK/ZacK filament-like structures were never observed to cross cell-cell borders and only traversed through not yet fully divided cells (Fig. 3B inlay and Fig. 3C). To further explore whether the ZicK/ZacK heteropolymer assembly is restricted to *Anabaena*, we proceeded to analyse ZicK and ZacK *in vivo* in an unrelated heterologous system using the *E. coli* split GFP assay (Wilson *et al.*, 2004). Clearly discernible filamentous-like structures (reminiscent of FilP-GFP (Bagchi *et al.*, 2008)) could be detected upon co-expression of ZicK and ZacK C-terminally fused to the split GFP products (ZicK-NGFP and ZacK-CGFP; Supplementary Fig. 5). This is in agreement with the lack of discernible structures upon expression of N-terminally GFP-fused ZicK and ZacK in *Anabaena* and is also in concert with the essential N-terminal domain for polymerization of IF and IF-like proteins (Heins and Aebi, 1994; Cabeen *et al.*, 2009; Cabeen *et al.*, 2011). Nonetheless, some indications for heteropolymerization were also present upon co-expression of NGFP-ZicK with ZacK-CGFP, which is in agreement with the BACTH results that indicated that both, N and C-terminal fusions of ZicK and ZacK are potentially able to interact with each other (Fig. 2B). The different heteropolymerization phenotype of ZicK/ZacK polymer in *E. coli* and *Anabaena* suggests that there are other so far unknown factors that modulate the specific ZicK/ZacK heteropolymerization phenotype, as shown in the following section.

**Fig. 3:**
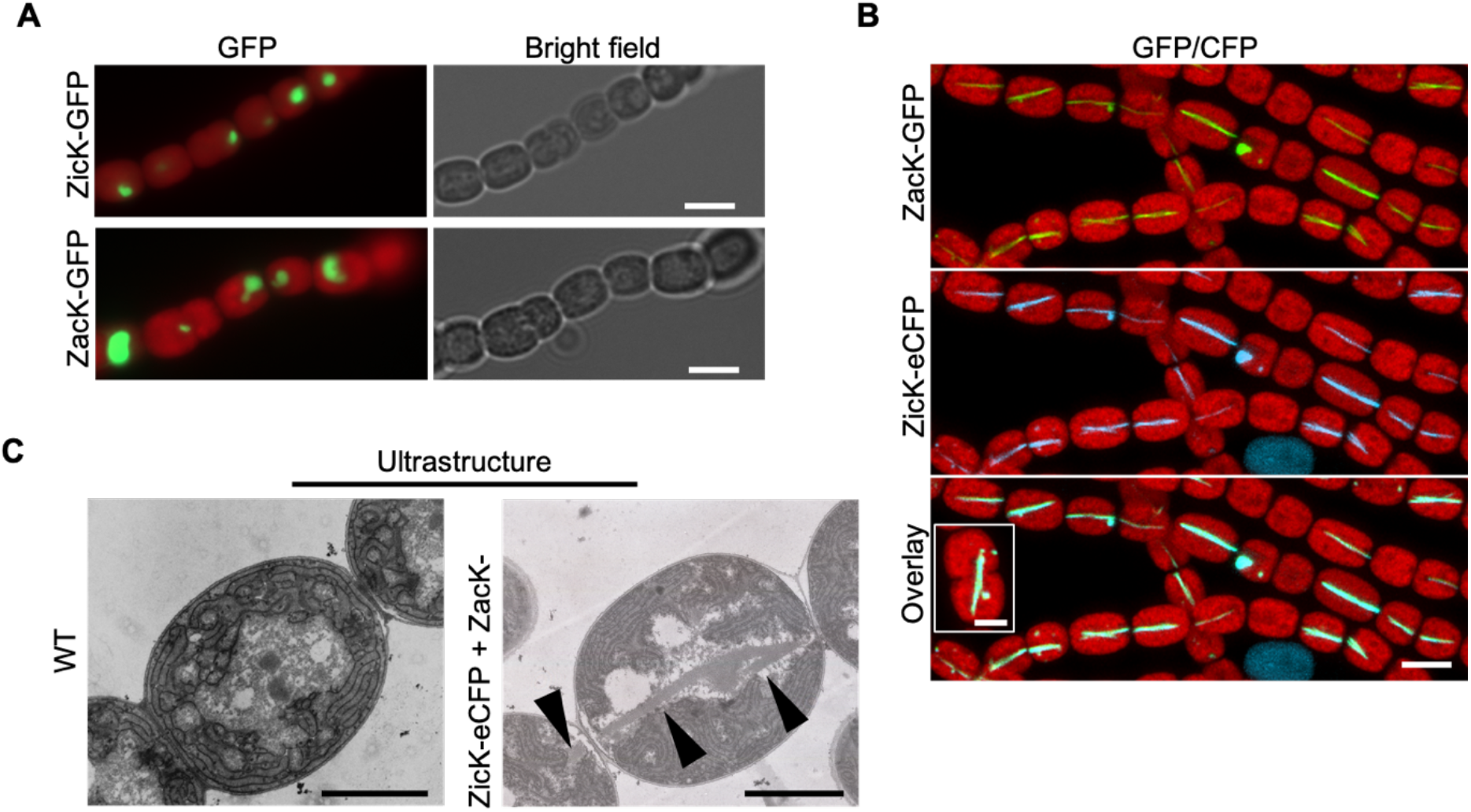
ZicK and ZacK form a heteropolymer *in vivo*. (**A**,**B**) Merged GFP or eCFP-fluorescence and chlorophyll autofluorescence (red) and bright field micrographs of *Anabaena* WT cells expressing (**A**) ZicK-GFP, ZacK-GFP or (**B**) co-expressing ZicK-eCFP and ZacK-GFP from P_petE_. (**B**) Inlay shows that ZicK/ZacK filaments only cross not yet fully divided cells. Scale bars: (A,B) 5 μm, (B inlay) 2.5 μm. (**C**) Electron micrographs of ultrathin sections of *Anabaena* WT and *Anabaena* cells co-expressing ZicK-eCFP and ZacK-GFP. Black arrows indicate electron-dense structures coinciding with the ZicK/ZacK heteropolymer observed in Fig. 4B. Scale bars: 1.6 μm.

### Deletion of *zicK* and *zacK* leads to defects in trichome and cell shape and *Anabaena* viability

In contrast to the obtained Δ*zicK*Δ*zacK* double mutant, single Δ*zicK* or Δ*zacK* mutant strains could not be generated, suggesting that the presence (or absence) of ZicK or ZacK alone is lethal for *Anabaena*. Further investigating the Δ*zicK*Δ*zacK* mutant phenotype revealed an altered trichome and cell shape phenotype and a reduced trichome viability (Fig. 4). Unlike the linear trichome growth pattern of the WT, the Δ*zicK*Δ*zacK* mutant strain grew as zigzagged trichomes (Fig. 4A), a phenotype that could be rescued by heterologous expression of P_zicK_::*zicK-zacK* from pRL25C but not from P_zicK_::*zicK-ecfp*+*zacK-gfp* (Supplementary Figs, 4C and 6A). Additionally, Δ*zicK*Δ*zacK* cells were significantly larger (WT: 27.42 ± 14.75 μm^3^; Δ*zicK*Δ*zacK*: 32.52 ± 12.54 μm^3^; P: <0.0001; Student’s t test) and significantly more round (WT: 0.8063 ± 0.1317; Δ*zicK*Δ*zacK*: 0.8530 ± 0.1130; P: <0.0001; Student’s t test) in comparison to the WT (Fig. 4B,C), reminiscent of the Δ*mreB* mutant strain (Hu *et al.*, 2007). The round and swollen cell phenotypes of the Δ*zicK*Δ*zacK* mutant strains are indicative of an impairment in cell wall integrity and/or defects in PG biogenesis as well as an elevated sensitivity to turgor pressure (Fenton *et al.*, 2016; Rojas and Huang, 2018). Consequently, we tested for an elevated sensitivity of the Δ*zicK*Δ*zacK* mutant to cell wall damaging enzymes. This showed that the Δ*zicK*Δ*zacK* mutant is slightly more sensitive to Proteinase K treatment but was unaffected by lysozyme treatment and still retained the ability to grow diazotrophically (*i.e*., on BG11_0_ plates) (Fig. 4D) and to form heterocysts (Supplementary Fig. 6B). More importantly, however, we found that the Δ*zicK*Δ*zacK* mutant lost the ability to grow in liquid culture (with and without agitation; Fig. 4E), which could be complemented with pRL25C carrying P_zicK_::*zicK-zacK* (Supplementary Fig. 6C), hinting for an elevated sensitivity to fluid shear stress or turgor pressure.

**Fig. 4:**
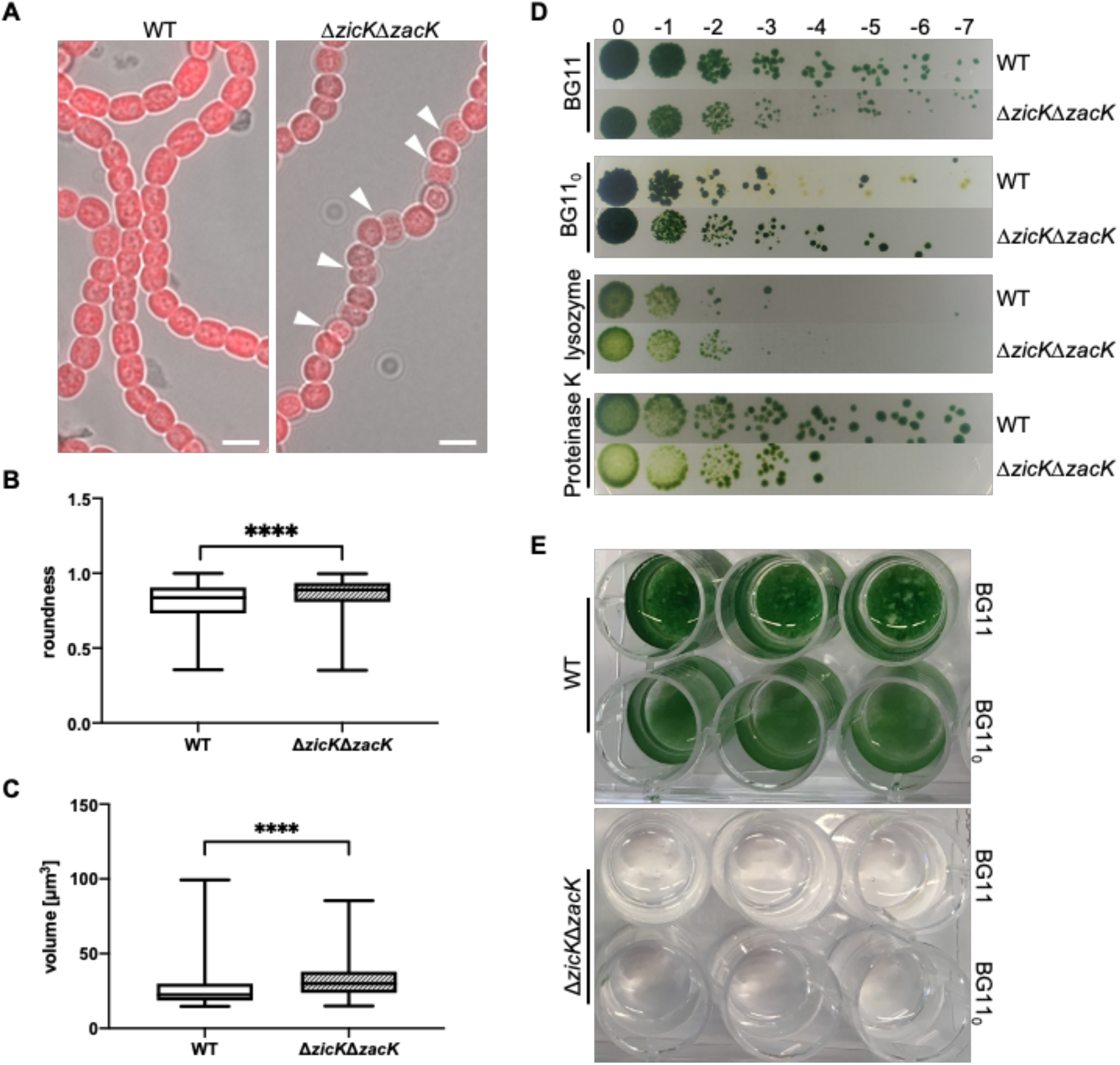
Deletion of *zicK* and *zacK alters trichome and cell shape as well as trichome viability*. (**A**) Merged chlorophyll autofluorescence and bright field micrographs of *Anabaena* WT and Δ*zicK*Δ*zacK* mutant grown on BG11 plates. White triangles indicate zigzagged trichome growth. Scale bars: 5 μm. (**B**) Cell roundness and (**C**) volume of *Anabaena* WT and Δ*zicK*Δ*zacK* mutant measured with Fiji imaging software A perfect circle is defined as roundness of 1. Error bars indicate standard deviations (*Anabaena* WT: n=537; Δ*zicK*Δ*zacK*: n=404). Values indicated with asterisks are significantly different from the WT. ****: p < .0001 (Student‘s t-test). (**D**) *Anabaena* WT and Δ*zicK*Δ*zacK* mutant were spotted onto BG11, BG11_0_ or BG11 plates supplemented with lysozyme or Proteinase K in triplicates of serial dilutions of factor 10 and grown until no further colonies arose in the highest dilution (n=2). (**E**) *Anabaena* WT and Δ*zicK*Δ*zacK* mutant were grown on BG11 plates, transferred to liquid BG11 and BG11_0_ medium and incubated for 12 d at standard growth conditions without shaking.

### ZicK and ZacK interact with proteins involved in cell shape and trichome integrity

Considering the impact of the deletion of *zicK* and *zacK* on cell and trichome shape and the assumed septal docking of the ZicK/ZacK heteropolymer, we next wanted to investigate whether both, ZicK and ZacK physically interact with other proteins known to function in cell shape determination and cell-cell communication. Using the BACTH assay, we found that both, ZicK and ZacK, interacted with the divisome-associated septal protein SepI (Springstein *et al.*, 2020a), the septal protein SepJ (Flores *et al.*, 2007), the cell shape-determining protein MreB as well an elongasome associated protein (ZipM; covered in more detail in subsequent study) and the *Anabaena* homolog to HmpF (here named HmpF_Ana_), whose homologs were shown to be involved in motility in *Nostoc punctiforme* ATCC 29133 (Cho *et al.*, 2017) and *Synechocystis* sp. PCC 6803 (Bhaya *et al.*, 2001; Springstein *et al.*, 2020b) (Fig. 5A). No interactions were found with FtsZ, FraC and FraD (Supplementary Fig. 7). We attempted to further confirm our interaction results with affinity co-elution experiments but found that Ni-NTA-bound ZicK and ZacK purified from *E. coli* readily precipitated upon transfer from denaturing to native buffer conditions, precluding further co-elution studies. Additionally, we observed that non-denaturing conditions failed to purify overexpressed CCRPs from *E. coli*, confirming their inherent insoluble nature, a property known to eukaryotic IFs (Kelemen, 2017). Instead, we surveyed for further interaction partners by anti-GFP co-immunoprecipitation experiments of *Anabaena* cells expressing ZicK-GFP and analysed co-precipitated proteins by LC-MS/MS analytics (all 27 identified possible interactors are listed in Supplementary File 3). This analysis confirmed that ZicK and ZacK interact with each other *in vivo* and further strengthened the observed association of ZicK with MreB (Fig. 5B). Furthermore, ZicK co-precipitated ParA (Fig. 5B), a walker A-type ATPase, involved in chromosome and plasmid partitioning (Lutkenhaus, 2012) and All4051 (also termed AnAKb), a protein associated with low-temperature resistance and potentially involved in cryoprotectant production (Ehira *et al.*, 2005).

**Fig. 5:**
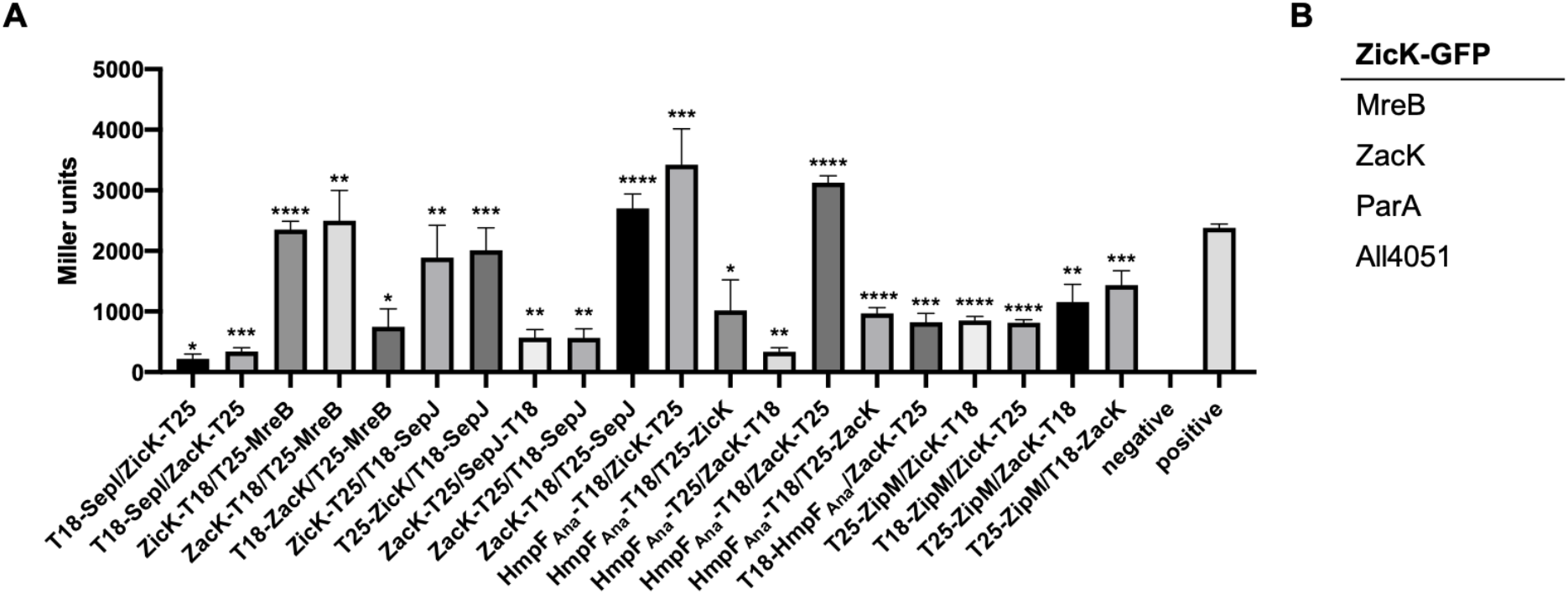
ZicK and ZacK interact with a multitude of *Anabaena* proteins involved in cell shape and multicellularity. (**A**) BACTH assays of *E. coli* cells co-expressing indicated T25 and T18 translational fusions of indicated pair-wise combinations of ZicK, ZacK, SepI, MreB, SepJ, HmpF_Ana_ and ZipM. Only translational fusion combinations that resulted in a significant interaction between two analysed proteins are shown. All other combinations were negative for interaction. Note that we used full-length proteins for all our BACTH analysis, including SepJ, whose precise subcellular localization remains to be identified (Ramos-León *et al.*, 2017; Springstein *et al.*, 2020a) but could be different in *E. coli* and *Anabaena* (Springstein *et al.*, 2020a). Quantitative values are given in Miller units, and the mean results from three independent colonies are presented. Negative: N-terminal T25 fusion construct of the respective protein co-transformed with empty pUT18C. Positive: Zip/Zip control. Error bars indicate standard deviations (n = 3). Values indicated with asterisks are significantly different from the negative control. *: p < .05, **: p < .005, ***: p < .0005, ****: p < .0001 (Student‘s t-test). (**B**) Excerpt of the identified specific interactors of ZicK-GFP. The full list is listed in Supplementary File 3.

### Deletion of *zicK* and *zacK* affects the localization of MreB and the chromosomes

As our BACTH analysis identified SepI and SepJ as interaction partners, and both proteins are involved in intercellular transport and cell-cell communication in *Anabaena* (Mullineaux *et al.*, 2008; Springstein *et al.*, 2020a), we proceeded to test whether the ΔzicKΔzacK mutant is also affected in solute diffusion using fluorescence recovery after photobleaching (FRAP) experiments of calcein stained ΔzicKΔzacK mutant. However, this analysis did not reveal any defect in cell-cell communication in the ΔzicKΔzacK mutant (Supplementary Fig. 8A-C) and hence ZicK and ZacK do not affect septal junction functionality. Additionally, electron microscopy of ultrathin sections of the ΔzicKΔzacK mutant, did not show any discernible differences in the ultrastructure of the cells compared to cells of *Anabaena* WT (Supplementary Fig. 8D). In accordance with a lack of interaction between FtsZ and ZicK/ZacK, FtsZ placement was unaffected in the ΔzicKΔzacK mutant as shown using anti-FtsZ immunofluorescence (Supplementary Fig, 9A). Following the lead of ZicK/ZacK interaction partners, we next analysed the localization of MreB in the *Anabaena* WT and the ΔzicKΔzacK mutant using a functional P_petE_::*gfp-*mreB fusion (Hu *et al.*, 2007). In *Anabaena* WT, we observed GFP-MreB filaments throughout the cells without any directional preferences and sometimes forming local foci (Fig. 6A). Even though GFP-MreB filaments were present in the Δ*zicK*Δ*zacK* mutant strain (Fig. 6A inlay), we only detected those filaments in non-rounded cells that seemingly had a WT-like phenotype (Fig 6A), accounting for 24% of counted cells (245 of 1040 cells counted), whereas in rounded/swollen cells of zigzagged trichomes, the GFP-MreB signals were restricted to the cell poles (Fig. 6A), accounting for 76% of counted cells (795 of 1040 counted cells). To further investigate the potential effect of *zicK* and *zacK* deletion on MreB and hence elongasome function, we stained sites of active cell wall biosynthesis using a fluorescent vancomycin derivate (Van-FL; (Daniel and Errington, 2003)). The staining pattern between the WT and the ΔzicKΔzacK mutant was indistinguishable but the fluorescence intensity levels were slightly decreased in the ΔzicKΔzacK mutant (Supplementary Fig. 9B,C). Nonetheless, this is likely accounted for by the reduced growth rate of the ΔzicKΔzacK mutant (Fig. 4D and general observation on growth plates).

**Fig. 6:**
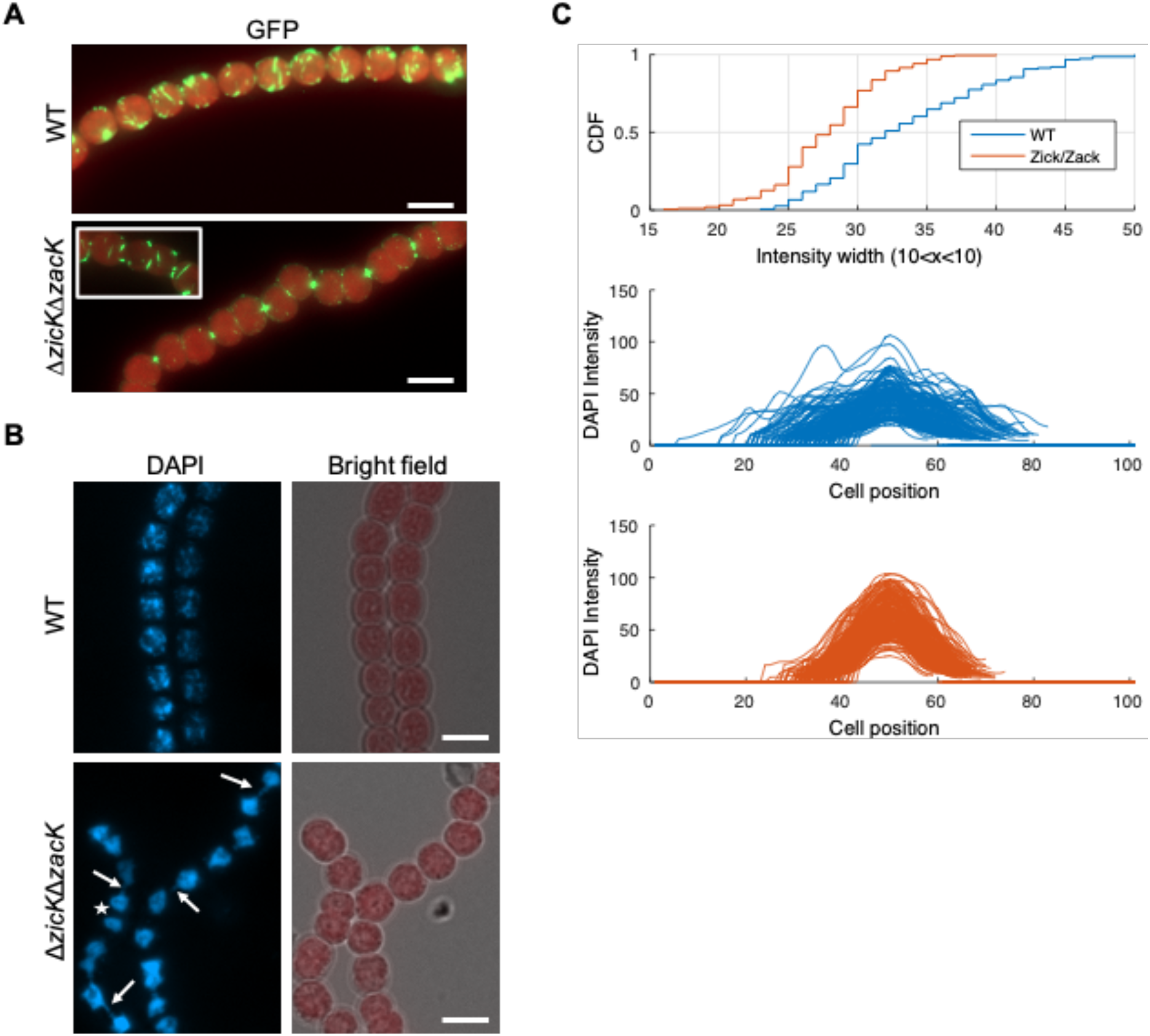
Effect of *zicK* and *zacK* deletion on MreB localization and DNA distribution. (**A**) Merged GFP fluorescence and chlorophyll autofluorescence micrographs of *Anabaena* WT and Δ*zicK*Δ*zacK* mutant expressing GFP-MreB from P_petE_. Cells were grown on BG11 plates. Scale bars: 5 μm. (**B**) DAPI fluorescence and merged bright field and chlorophyll autofluorescence micrographs of *Anabaena* WT and Δ*zicK*Δ*zacK* mutant on BG11 plates. White arrows indicate strings of DNA that seemingly traverse from one cell to the other. Notably, no such strings are observed in dividing cells (white star), suggesting that it is an effect that occurs after complete cell division. Although, we note that high resolution microscopy would be needed to fully resolve this observation. Scale bars: 5 μm. (**C**) Plot profile showing a cumulative distribution function (CDF) of the DAPI signal intensities of pixels (grey value) along *Anabaena* WT and Δ*zicK*Δ*zacK* mutant cells (n=151 for each strain) in arbitrary units (a.u.) and arranged to the respective peak maxima. The focal area size in the Δ*zicK*Δ*zacK* mutant was smaller in comparison to the *Anabaena* WT (P=6.8×10^−117^, using Wilcoxon test). Notably, the comparison of cell lenght among the strains reveals a similar result: the Δ*zicK*Δ*zacK* mutant cell size was smaller in comparison to the *Anabaena* WT (P=7.2×10^−117^, using Wilcoxon test). Consequently, we compared the area of the focal DAPI staining decided by the cell size among the strains. This, however, revealed that this ratio is not significantly different between the Δ*zicK*Δ*zacK* mutant and the *Anabaena* WT.

Considering the interaction of ZicK/ZacK with ParA, we further tested for a function of ZicK and ZacK in DNA placement and compared the DNA distribution in the WT and the ΔzicKΔzacK mutant as measured by distribution of 4′,6-Diamidin-2-phenylindol (DAPI) staining intensity (Fig. 6B,C). For that, we calculated the width of the DAPI focal area as the range of DAPI staining around the maximum intensity focus (±10 grey intensity in arbitrary units). This revealed that the staining focal area size was significantly different among the WT and the ΔzicKΔzacK mutant. The DAPI signal observed in the Δ*zicK*Δ*zacK* mutant appears more condensed, and indeed, the Δ*zicK*Δ*zacK* mutant focal DAPI area was smaller than the WT (Fig. 6C). Unlike the WT, DAPI signals in the Δ*zicK*Δ*zacK* mutant was also observed between two neighbouring cells (Fig. 6B). Overall our results suggest the involvement of ZicK/ZacK in DNA distribution and segregation in dividing cells.

## Discussion

Here we provide evidence for the capacity of two *Anabaena* CCRPs, which we termed ZicK and ZacK, to form polymers *in vitro* and *in vivo*. While the previously described prokaryotic filament-forming CCRPs formed homopolymers (Ausmees *et al.*, 2003; Yang *et al.*, 2004; Bagchi *et al.*, 2008; Specht *et al.*, 2011), ZicK and ZacK exclusively assembled into a heteropolymer *in vitro* and *in vivo*, thus revealing a new property of bacterial CCRPs. The inherent heteropolymerization tendency of ZicK and ZacK was confirmed in a heterologous and evolutionary distant *E. coli* system, which was previously used to investigate other known CCRPs such as Scc from *Leptospira biflexa* (England *et al.*, 2005) or Crescentin (Ingerson-Mahar *et al.*, 2010). Although heteropolymerization has previously been described for prokaryotic cytoskeletal proteins, none of those polymerization pairs both belonged to the group of CCRPs. BacA and BacB, members of the widely conserved class of bactofilins, both independently polymerize into filaments *in vitro*, co-localize *in vivo* in *C. crescentus* and interact directly with each other as indicated by co-immunoprecipitation analysis (Kühn *et al.*, 2010). Unlike CCRPs, whose self-interaction is based on the high degree of CC domains, in bactofilins, the DUF583 domain is proposed to mediate protein polymerization (Kühn *et al.*, 2010). Despite compelling evidence for co-assembly and shared functional properties, heteropolymerization of BacA and BacB hasn’t been studied *in vitro*. Another interesting pair of potential co-polymerizing cytoskeletal proteins that both independently assemble into homopolymers but also co-align *in vivo* and affect each other’s properties are Crescentin and the CtpS enzyme from *C. crescentus* (Ingerson-Mahar *et al.*, 2010). Although, again, the co-assembly *in vitro* is not reported in the literature.

Despite the numerous independently confirmed heteropolymerization properties of ZicK and ZacK, we note, however, that the results from our *in vivo* experiments are based on artificial expression of the two CCRPs. We hypothesize that the absence of a ZicK/ZacK heteropolymer in strains expressing ZicK-GFP or ZacK-GFP alone (with the WT *zicK* and *zacK* alleles still present) may be due to a dosage-dependent effect, where the presence of unequal concentration of ZicK or ZacK in the cell leads to protein aggregates. Our observation of ZicK-GFP or ZacK-GFP aggregates when they were expressed alone in the Δ*zicK*Δ*zacK* mutant strain supports the dosage effect hypothesis. Also, in our *in vitro* polymerization assay, ZicK and ZacK only formed clear and distinct filament-like structures when both proteins are present in equal concentrations. Nonetheless, co-expressed of ZicK-eCFP and ZacK-GFP were not able to complement the ΔzicKΔzacK mutant. Attempts to express both proteins fused to a fluorophore from the native promoter remained unsuccessful, possibly a result of the close genomic proximity. Furthermore, the genomic neighbourhood of *zicK* and *zacK* suggests that the ZicK/ZacK heteropolymer formation could be relying on co-translational assembly (e.g., as observed for LuxA/LuxB (Shieh *et al.*, 2015)). Co-translational assembly of natively present ZicK and ZacK would lead to an efficient binding of the two subunits such that the additional expression of one unit only in excess (*i.e*., ZicK-GFP or ZacK-GFP alone) would lead to the formation of aggregates. As such, it remains to be elucidated to what extent the ZicK/ZacK heteropolymer exists in *Anabaena*. Although, we could not identify any protein filaments in our ultrathin sections from *Anabaena* WT, other studies have previously described filamentous strings and even longitudinal cell-spanning polymers in multicellular *Anabaena* and *Nostoc* strains (Jensen and Ayala, 1980; Bermudes *et al.*, 1994). Despite compelling evidence for the existence of a cyanobacterial Z-ring structure during cell division (Sakr *et al.*, 2006b; Sakr *et al.*, 2006a; Ramos-León *et al.*, 2015; MacCready *et al.*, 2017; Corrales-Guerrero *et al.*, 2018; Camargo *et al.*, 2019), no Z-ring ultrastructures have yet been identified and consequently, the absence of longitudinal ZicK/ZacK filaments in ultrathin sections does not rule out that they exist but could rather indicate that they could not be visualized yet.

Our results indicate that ZicK and ZacK are associated with the elongasome (through their interaction with MreB) and proteins in the septal cell wall (through the interaction with SepJ and SepI) and affect cellular DNA placement (Fig. 7). A function of ZicK/ZacK in chromosome segregation, would be in concert with the identified interaction of ZicK with ParA, this, however, remains to be elucidated as it could also be an indirect consequence of the swollen/rounded cell shape in the Δ*zick*Δ*zack* mutant. Nonetheless, so far no chromosome partitioning system has yet been identified in multicellular cyanobacteria (Hu *et al.*, 2007). In *E. coli*, *B. subtilis* and *C. crescentus*, MreB functions in chromosome segregation while deletion of *mreB* did not affect chromosome segregation in *Anabaena* but induced a swollen cell phenotype (Hu *et al.*, 2007), similar to the Δ*zick*Δ*zack* mutant. Consequently, MreB and ZicK/ZacK likely share functional properties but are not exclusively involved in the same cellular processes. Swollen cell morphotypes were also described for *Anabaena* or *Synechocystis* mutants lacking penicillin binding proteins (PBPs), which are enzymes that are directly involved in cell wall biogenesis through the modification of the PG layer (Lázaro *et al.*, 2001; Leganés *et al.*, 2005; Burnat *et al.*, 2014). This presumed link of ZicK/ZacK to the actin-like MreB cytoskeleton and the PG biogenesis apparatus is also indicated by the altered localization of GFP-MreB and the decreased PG staining intensity in the Δ*zicK*Δ*zacK* mutant strain. Consequently, ZicK and ZacK might indirectly act to positively regulate PG biogenesis, although, we cannot exclude that the reduced staining intensity in the Δ*zicK*Δ*zacK* mutant simply reflects the slower growth rate of this strain. An interaction or involvement of prokaryotic filament-forming CCRPs with MreB and PG synthesis were previously observed in other bacteria. Examples are the gliding motility in *Myxococcus xanthus,* where a multiprotein complex, including the filament-forming CCRP AglZ and MreB, were found to coordinate type A-motility (Schumacher and Søgaard-Andersen, 2017). Similarly, the curved morphotype of *C. crescentus* is induced by Crescentin, which functionally associates with MreB and likely modulates PG biogenesis by exuding local mechanical forces to the cell membrane (Charbon *et al.*, 2009; Lin and Thanbichler, 2013). Other aspects like a decreased cell envelope permeability of the Δ*zicK*Δ*zacK* mutant are also conceivable, although we did not detect any cell wall defects in the Δ*zicK*Δ*zacK* mutant. MreB and the elongasome are the main determinants of the PG exoskeleton, which provides the cell with structural integrity and resistance to turgor pressure (Typas *et al.*, 2012). The lack of liquid growth of the Δ*zicK*Δ*zacK* mutant would also argue for a defect in the resistance to turgor pressure.

**Fig. 7:**
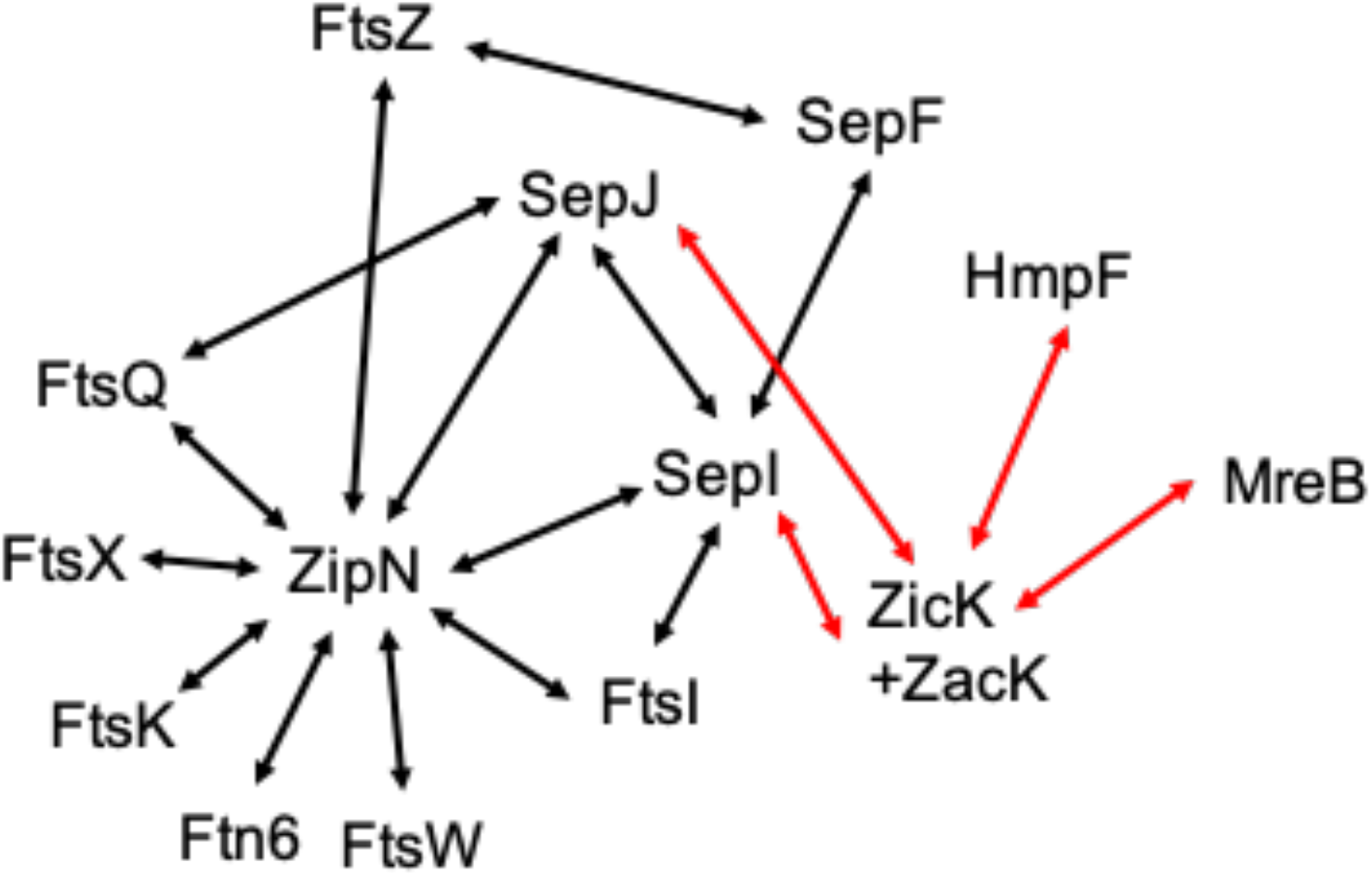
Interaction network of known divisome, elongasome and septal proteins in *Anabaena*. A model for a partial divisome, elongasome and septal junction network in *Anabaena* as deduced from BACTH and co-IP analyses. Black arrows indicate interactions that have been previously described by (Ramos-León *et al.*, 2015; Camargo *et al.*, 2019; Springstein *et al.*, 2020a). Red arrows indicate interactions identified in the current analysis.

Together with the cell shape-determining protein MreB, ZicK and ZacK could possibly contribute to normal cell shape and relay trichome shape-stabilizing properties to neighbouring cells in the trichome by means of their association with the filament stabilizing protein SepJ (Fig. 7). As such, they are important for maintaining the linear *Anabaena* trichome phenotype. ZicK and ZacK polymers might constitute stabilizing platforms or scaffolds for other proteinaceous structures, similarly to the stabilizing function of the eukaryotic cytoskeleton for cell-cell contacts (*i.e*., desmosomes). Furthermore, ZicK shares *in silico* predicted structural similarities with the spectrin repeats of plectin, a well-described eukaryotic cytolinker protein. Plectin link the three eukaryotic cytoskeletal systems (actin filaments, microtubules and IFs), thereby contributing to the resistance to deformation of vertebrate cells (Alberts *et al.*, 2014). They stabilize desmosomes and are hence directly involved in cell-cell contact integrity (Leung *et al.*, 2002). An analogous cytolinker function of ZicK could explain why ZacK alone did not form properly folded protein filaments on its own and suggests that ZacK requires ZicK as the linking protein for polymerization. As plectin not only stabilizes but also dynamically disassembles IF protein filaments (*i.e*., vimentin) in a concentration-dependent manner (Birchler *et al.*, 2001), this would further support a dosage-dependent effect of ZicK and ZacK for heteropolymerization.

The conserved combination of ZicK and ZacK in heterocystous cyanobacteria that form linear trichomes (Fig. 1C, Supplementary File 2) highlights a potential function of ZicK and ZacK for the maintenance of the linear trichome. The Δ*zicK*Δ*zacK* mutant had a zigzagged phenotype and was unable to grow in liquid culture. We hypothesize that the loss of trichome linear shape led to an increase in accessible surface for the acting mechanical forces in liquid (Persat *et al.*, 2015), including fluid shear stress (Park *et al.*, 2011), ultimately resulting in forces that cannot be endured by the abnormal mutant trichomes. The loss of ZicK and/or ZacK in heterocystous cyanobacteria species that are in symbiosis (e.g., *N. azollae*) or form true-branching or multiseriate filaments (e.g., *Fischerella* or *Chlorogloeopsis*, respectively) may suggest that these species are less sensitive to mechanical stress (i.e., due to their interaction with the host or complex filament formation). The key hallmarks of permanent bacterial multicellularity are morphological differentiation and a well-defined and reproducible shape, termed patterned multicellularity (Claessen *et al.*, 2014). Besides the highly reproducible cell division, proliferation and cell differentiation in sporulating actinomycetes (Flärdh *et al.*, 2012), the reproducible linear trichomes in filamentous cyanobacteria are considered a major contributor to the cyanobacterial patterned multicellularity (Claessen *et al.*, 2014; Herrero *et al.*, 2016), manifesting a selective advantage to biotic and abiotic environmental factors (Young, 2006; Singh and Montgomery, 2011). Our results indicate that ZicK and ZacK serve as regulators of the typical linear *Anabaena* trichome and as such as regulators of *Anabaena* patterned multicellularity. The evolution of patterned multicellularity is considered an important step towards a sustainable division of labour and the development of cell differentiation (Claessen *et al.*, 2014). Our study provides initial evidence for a role of two heteropolymer-forming CCRPs in the evolution and maintenance of cyanobacterial multicellular forms.

## Acknowledgments

We thank Myriam Barz, Katrin Schumann, Lisa-Marie Philipp, Lisa Stuckenschneider and Claudia Menzel for their assistance in the experimental work and Andreas Tholey for help with the mass spectrometry analysis. FRAP experiments were performed at the Facility for Imaging by Light Microscopy (FILM) at Imperial College London. This study was supported by the German science foundation (DFG) (Grant No. STU513/2-1) and a Fondecyt Grant (Grant No. 11170842), both awarded to KS. IM was supported by German science foundation (DFG) (Grant SFB766). DJN was supported by the BBSRC as part of the joint NSF Ideas Lab grant on ‘Nitrogen: improving on nature’ (Grant No. BB/L011506/1).

## Author contribution

BLS and KS designed the study. BLS established and performed the experimental work with contributions from MLT and JW. CW and TD performed comparative genomics analysis. DJN performed FRAP assays and IM carried out ultratstructure analyses. AOH and AT analysed protein samples by mass spectrometry. BLS, TD and KS drafted the manuscript with contributions from all co-authors.

## Data availability

All data generated or analysed during this study are included in this published article (and its supplementary files).

## Material and methods

### Bacterial strains and growth conditions

*Anabaena* sp. PCC 7120 was obtained from the Pasteur Culture Collection (PCC) of cyanobacteria (France). Cells were grown photoautotrophically in BG11 or without combined nitrogen (BG11_0_) at constant light with a light intensity of 30 μmol m^−2^ s^−1^ in liquid culture or on agar plates (1.5% w/v agar). When appropriate, 5 μg ml^−1^ spectinomycin (Sp), 5 μg ml^−1^ streptomycin (Sm) or 30 μg ml^−1^ neomycin (Nm) was added to strains carrying respective plasmids or chromosomal insertions. In some cases, basal copper-regulated *petE*-driven expression of gene candidates in *Anabaena* cells was lethal or growth inhibiting, therefore these strains were grown in BG11 without copper and protein expression was later induced by the addition of CuSO_4_ at indicated concentrations to the culture. *E. coli* strains DH5α, DH5αMCR, XL1-blue and HB101 were used for cloning and conjugation by triparental mating. BTH101 was used for BACTH system and BL21 (DE3) was used for expression of His_6_-tagged proteins in *E. coli*. All *E. coli* strains were grown in LB medium containing the appropriate antibiotics at standard concentrations. Supplementary Tables 1-4 list all used bacterial strains, plasmids and oligonucleotides.

### Prediction of coiled-coil-rich proteins

Genome sequence of *Anabaena* (GCA_000009705.1) was analysed by the COILS algorithm (Lupas *et al.*, 1991) as previously described (Bagchi *et al.*, 2008). The algorithm was run with a window width of 21 and the cut-off for amino acids in coiled-coil conformation was set to ≥80 amino acid residues. The resulting set of protein candidates was further manually examined with online available bioinformatic tools, including NCBI Conserved Domain Search (Boratyn *et al.*, 2012), NCBI BLAST (Altschul *et al.*, 1990), TMHMM (Sonnhammer *et al.*, 1998) and I-TASSER (Zhang, 2009). Protein candidates exhibiting BLAST hits involved in cytoskeletal processes or similar domain architectures as known IF and IF-like proteins like Crescentin, FilP, vimentin, desmin or keratin were selected, and enzymatic proteins as well as proteins predicted to be involved in other cellular processes were excluded.

### Distribution of homologs in cyanobacteria

Homologs to the *Anabaena* proteins were extracted from pre-calculated cyanobacterial protein families (Springstein *et al.*, 2020b). Conserved syntenic blocks (*i.e*., gene order) were identified using CSBFinder-S (Svetlitsky *et al.*, 2020).

### RNA isolation and cDNA synthesis

RNA from *Anabaena* WT was isolated using the Direct-zol™ RNA MiniPrep Kit (Zymo Research) according to the manufacturer’s instructions. RNA was isolated in technical triplicates from 10 ml cultures. Isolated RNA was treated with DNA-free™ Kit (2 units rDNAs/reaction; Thermo Fischer Scientific) and 200 ng RNA was reverse transcribed using the qScript™ cDNA Synthesis Kit (Quanta Biosciences). RT-PCR of cDNA samples for *rnpB*, *zicK*, *zacK* and *zicK*+*zacK* was performed using primer pairs #1/#2, #3/#4, #5/#6 and #3/#8, respectively.

### Transformation

*Anabaena* was transformed by triparental mating as previously described (Ungerer and Pakrasi, 2016). Briefly, 100 μl of overnight cultures of DH5α carrying the conjugal plasmid pRL443 and DH5αMCR carrying the cargo plasmid and the helper plasmid pRL623, encoding for three methylases, were mixed with 200 μl *Anabaena* culture (for transformation into the *ΔzickΔ*zacK mutant, cells were scraped from the plate and resuspended in 200 μl BG11). This mixture was directly applied onto sterilized nitrocellulose membranes (Amersham Protran 0.45 NC) placed on top of BG11 plates supplemented with 5% (v/v) LB medium. Cells were incubated in the dark at 30 °C for 6-8 h with subsequent transfer of the membranes to BG11 plates and plates were placed under standard growth conditions. After 24 h, membranes were transferred to BG11 plates supplemented with appropriate antibiotics.

### Plasmid construction

Ectopic expression of *Anabaena* protein candidates was achieved from a self-replicating plasmid (pRL25C (Wolk *et al.*, 1988)) under the control of the copper-inducible *petE* promoter (P_petE_) or the native promoter (predicted by BPROM (Solovyev and Salamov, 2011)) of the respective gene. All constructs were verified by Sanger sequencing (Eurofins Genomics). Plasmids were created using standard restriction enzyme-based techniques or Gibson assembly. Information about precise plasmid construction strategies are available from the authors upon request.

### Anabaena mutant strain construction

The *ΔzickΔ*zacK mutant strain was generated using the pRL278-based double homologous recombination system employing the conditionally lethal *sacB* gene (Cai and Wolk, 1990). For this, 1500 bp upstream and downstream of *zick-zacK were* generated by PCR from *Anabaena* gDNA. Upstream region of *zicK* was amplified using primers #97/#98 and downstream region of *zacK* was amplified using primers #99/#100. The respective upstream and downstream homology regions flanking the CS.3 cassette (amplified with primer #95/#96 from pCSEL24) were then inserted into PCR-amplified pRL278 (using primer #93/#94) by Gibson assembly, yielding pTHS166. *Anabaena* transformed with pTHS166 plasmids was subjected to several rounds of re-streaking on new plates (about 5-8 rounds). To test for fully segregated clones, colony PCRs were performed. For this, *Anabaena* cells were resuspended in 10 μl sterile H2O of which 1 μl was used for standard PCR with internal *zicK* and *zacK* gene primers #3/#6. Correct placement of the CS.3 cassette was then further confirmed using CS.3 cassette primers with binding sites outside of the 5’ and 3’ flanks used for homologous recombination (primers #95/#102 and #101/#96).

### Fluorescence microscopy

Bacterial strains grown in liquid culture were either directly applied to a microscope slide or previously immobilized on a 2% (w/v) low-melting agarose in PBS (10 mM Na_2_HPO_4_, 140 mM NaCl, 2.7 mM KCl, 1.8 mM KH_2_PO_4_, pH 7.4) agarose pad and air dried before microscopic analysis. Epifluorescence was done using an Axio Imager.M2 light microscope (Carl Zeiss) equipped with Plan-Apochromat 63x/1.40 Oil M27 objective and the AxioCam MR R3 imaging device (Carl Zeiss). GFP, Alexa Fluor 488 and BODIPY™ FL Vancomycin (Van-FL) fluorescence was visualized using filter set 38 (Carl Zeiss; excitation: 470/40 nm band pass (BP) filter; emission: 525/50 nm BP). Chlorophyll auto-fluorescence was recorded using filter set 15 (Carl Zeiss; excitation: 546/12 nm BP; emission: 590 nm long pass). When applicable, cells were previously incubated in the dark at RT for about 5 min with 10 μg ml^−1^ DAPI (final concentration) to stain intracellular DNA. For visualization of DAPI fluorescence filter set 49 (Carl Zeiss; excitation: G 365 nm; emission: 455/50 nm) was employed. For confocal laser scanning microscopy, the LSM 880 Axio Imager 2 equipped with a C-Apochromat 63x/1.2 W Korr M27 objective and an Airyscan detector (Carl Zeiss) was used and visualization of GFP, eCFP and chlorophyll auto-fluorescence was done using Zen black smart setup settings.

### Transmission electron microscopy

For ultra-structure analysis, *Anabaena* trichomes were fixed with 2.5% (v/v) glutaraldehyde, immobilized in 2% (w/v) agarose, treated with 2% (v/v) potassium permanganate and dehydrated through a graded ethanol series (Mohr *et al.*, 2010). The fixed cells were infiltrated by ethanol:EPON (2:1 to 1:2 ratio) and embedded in pure EPON. Ultrathin sections were prepared with a Leica UC6i Ultramicrotome, transferred to formvar coated copper grids (Science Services GmbH München) and post-stained with uranyl acetate and lead citrate (Fiedler *et al.*, 1998). Micrographs were recorded at a Philips Tecnai10 electron microscope at 80 kV.

### Calcein labelling and fluorescence recovery after photobleaching (FRAP) experiments

For FRAP experiments, *Anabaena* WT and *ΔzickΔzacK* mutant strain were grown on BG11 plates, resuspended in BG11 liquid media and washed three times in 1 ml BG11 (3,000 × *g*, 5 min). Cells were then resuspended in 0.5 ml BG11 and incubated with 10 μl calcein-AM (Cayman Chemical, 1 mg ml^−1^ in DMSO) for 1 h at 30 °C in the dark. To remove excess staining solution the cells were washed four times with 1 ml BG11. Subsequently, the cells were spotted on BG11 agar for visualization by confocal laser scanning microscopy (Leica TCS SP5; HCX PL APO 63x 1.40-0.60 OIL CS). Calcein was excited at 488 nm and fluorescence emission monitored in the range from 500 to 530 nm with a maximally opened pinhole (600 μm). FRAP experiments were carried out by an automated routine as previously described (Mullineaux *et al.*, 2008). After recording an initial image, selected cells were bleached by increasing the laser intensity by a factor of 5 for two subsequent scans and the fluorescence recovery followed in 0.5 s intervals for 30 s was recorded using the Leica LAS X software. Exchange coefficients (*E*) were then calculated as previously described (Mullineaux *et al.*, 2008; Nieves-Morión *et al.*, 2017).

### BODIPY™ FL Vancomycin (Van-FL) staining

Van-FL staining of strains grown on BG11 plates was essentially performed as described previously (Lehner *et al.*, 2013; Rudolf *et al.*, 2015). Briefly, cells were resuspended in BG11 medium, washed once in BG11 by centrifugation (6500 × *g*, 4 min, RT) and incubated with 5 μg ml^−1^ Van-FL (dissolved in methanol; Thermo Fischer Scientific). Cells were incubated in the dark for 1 hour at 30 °C, washed three times with BG11 and immobilized on an agarose pad. Van-FL fluorescence signals were then visualized using epifluorescence microscopy with an excitation time of 130 ms. Arithmetic mean fluorescence intensities were recorded from the septa between two cells with a measured area of 3.52 μm^2^ using the histogram option of the Zen blue 2.3 software (Carl Zeiss).

### Alcian blue staining

*Anabaena* WT and ΔzicKΔzicK cells were grown on BG11_0_ plates, re-suspended in BG11_0_ liquid medium and stained with 0.05% (w/v) alcian blue (final concentration). Polysaccharide staining of cells immobilized on an agarose pad was observed with an Axiocam ERc 5s color camera (Carl Zeiss).

### Data analysis

Cell volume and roundness were determined using the imaging software ImageJ (Schneider *et al.*, 2012), a perfect circle is defined to have a roundness of 1. Cell volume was calculated based on the assumption of an elliptic cell shape of *Anabaena* cells using the Major Axis and Minor Axis values given by ImageJ and the formula for the volume of an ellipsoid

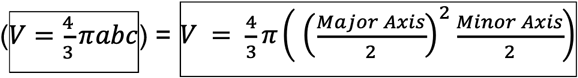

Distribution of DAPI fluorescence signal intensity was analysed in ImageJ with the Plot Profile option along 151 single cells with the rectangle tool. The resulting grey values were arranged according to the maximum intensity focus and the width of the DAPI focal area was calculated as the range of DAPI staining around the maximum (±10 grey value in arbitrary units). Statistical tests were performed with MatLab© (MathWorks) or GraphPad Prism v.8.

### Bacterial two-hybrid and beta galactosidase assays

Chemically competent *E. coli* BTH101 cells were co-transformed with 5 ng of plasmids carrying the respective T18 and T25 translational fusion constructs, plated onto LB plates supplemented with 200 μg ml^−1^ X-gal, 0.5 mM IPTG, Amp, Km and grown at 30°C for 24-36 h. Interactions were quantified by beta-galactosidase assays from three independent colonies. For this aim, cultures were grown for two days at 20 °C in LB Amp, Km, 0.5 mM IPTG and beta-galactosidase activity was recorded as described in the manufacturer’s instructions (Euromedex; BACTH System Kit Bacterial Adenylate Cyclase Two-Hybrid System Kit) in a 96 well plate format as previously described (Karimova *et al.*, 2012).

### GFP-fragment reassembly assay

Chemically competent *E. coli* BL21 (DE3) were co-transformed with indicated plasmid combinations, plated on LB Amp, Km and grown over night at 37 °C. Liquid overnight cultures of single colonies of the respective plasmid-bearing *E. coli* strains were then diluted 1:40 in the same medium the following day. Cells were grown for 2 h at 37 °C, briefly acclimated to 20 °C for 10 min and protein expression was induced with 0.05 mM IPTG and 0.2% (w/v) L-arabinose. Pictures of induced cultures grown at 20 °C were taken 48 h after induction.

### Co-immunoprecipitation

About 20-30 ml of the respective *Anabaena* cultures were pelleted by centrifugation (4800 × *g*, 10 min, RT), cells were washed twice by centrifugation (4800 × *g*, 10 min, RT) with 40 ml PBS and then resuspended in 1 ml lysis buffer (PBS-N: PBS with 1% (v/v) NP-40) supplemented with protease inhibitor cocktail (PIC; cOmplete™, EDTA-free Protease Inhibitor Cocktail, Sigma-Aldrich). Cells were lysed using the VK05 lysis kit (Bertin) in a Precellys® 24 homogenizer (3 strokes for 30 s at 6500 rpm) and cell debris was pelleted by centrifugation (30 min, 21,100 × *g*, 4 °C). 50 μl μMACS anti-GFP MicroBeads (Miltenyi Biotec) were added to the resulting cell-free supernatant and incubated for 1 h at 4 °C with mild rotation. Subsequently, the sample was loaded onto μColumns (Miltenyl Biotec), washed two times with 1 ml lysis buffer and eluted in 50 μl Elution Buffer (50 mM Tris HCl pH 6.8, 50 mM DTT, 1% (w/v) SDS, 1 mM EDTA, 0.005% (w/v) bromophenol blue, 10% (v/v) glycerol; Miltenyl Biotec). Samples were stored at −80 °C until further use.

### Mass spectrometry

Mass spectrometry of co-precipitated proteins was performed as previously described (Springstein *et al.*, 2020a).

### Immunofluorescence

Immunolocalization of FtsZ in *Anabaena* WT and Δ*zicK*Δ*zacK* mutant was essentially performed as previously described (Ramos-León *et al.*, 2015). For this, strains were scraped off from growth plates (BG11 and BG11_0_ plates), resuspended in a small volume of distilled water and air-dried on Polysine^®^ adhesion slides (Menzel) at RT followed by fixation and permeabilization with 70% ethanol for 30 min at −20 °C. Cells were allowed to air dry for 30 min at RT and then washed two times with PBST (PBS supplemented with 0.1% (v/v) Tween-20) for 2 min. Unspecific binding sites were blocked for 30 min at RT with blocking buffer (1x Roti®-ImmunoBlock in PBST; Carl Roth) and afterwards rabbit anti-FtsZ (Agrisera; raised against *Anabaena* FtsZ; 1:150 diluted) antibody in blocking buffer was added to the cells and incubated for 1.5 h at RT in a self-made humidity chamber followed by five washing steps with PBST. 7.5 μg ml^−1^ (final concentration) Alexa Fluor 488-conjugated goat anti-rabbit IgG (H+L) secondary antibody (Thermo Fischer Scientific) in blocking buffer was added to the cells and incubated for 1 h at RT in the dark in a self-made humidity chamber. Subsequently, cells were washed five times with PBST, air dried and mounted with ProLong™ Diamond Antifade Mountant (Thermo Fischer Scientific) overnight at 4 °C. Immunolocalization of FtsZ was then analysed by epifluorescence microscopy.

### Spot assays

For spot assays, *Anabaena* WT and Δ*zicK*Δ*zacK* mutant strain were grown on BG11 growth plates, resuspended in BG11 liquid medium and adjusted to an OD_750_ of 0.4. Cells were then spotted in triplicates of 5 μl onto the respective growth plates containing either no additives (BG11 or BG11_0_), 50 μg ml^−1^ Proteinase K or 100 μg ml^−1^ lysozyme in serial 1/10 dilutions and incubated under standard growth conditions until no further colonies arose in the highest dilution.

### Protein purification and in vitro filamentation assays

For protein purification, *E. coli* BL21 (DE3) cells carrying His-tagged protein candidates were grown in overnight cultures at 37 °C and 250 rpm. The next day, overnight cultures were diluted 1:40 in the same medium and grown at 37 °C until they reached an OD_600_ of 0.5-0.6. Protein expression was induced with 0.5 mM IPTG for 3-4 h at 37 °C and 250 rpm. Afterwards, cell suspensions of 50 ml aliquots were harvested by centrifugation, washed once in PBS and stored at −80 °C until further use. For *in vitro* filamentation assays, cell pellets were resuspended in urea lysis buffer (ULB: 50 mM NaH_2_PO_4_, 300 mM NaCl, 25 mM imidazole, 6 M urea; pH 8.0) and lysed in a Precellys® 24 homogenizer (3x 6500 rpm for 30 s) using the 2 ml microorganism lysis kit (VK01; Bertin) or self-packed Precellys tubes with 0.1 mm glass beads. The resulting cell debris was pelleted by centrifugation at 21,000 × *g* (10 min, 4 °C) and the supernatant was incubated with 1 ml HisPur™ Ni-NTA resin (Thermo Fischer Scientific) for 1 h at 4°C in an overhead rotator. The resin was washed five times with 4x resin-bed volumes ULB and eluted in urea elution buffer (UEB: ULB supplemented with 225 mM imidazole). Total protein concentration was measured using the Qubit® 3.0 Fluorometer (Thermo Fischer Scientific). Filament formation of purified proteins was induced by overnight dialysis against polymerization buffer (PLB: 50 mM PIPES, 100 mM KCl, pH 7.0; HLB: 25 mM HEPES, 150 mM NaCl, pH 7.4; or 25 mM HEPES pH 7.5) at 20 °C and 180 rpm with three bath changes using a Slide-A-Lyzer™ MINI Dialysis Device (10K MWCO, 0.5 ml or 2 ml; Thermo Fischer Scientific). Purified proteins were stained with an excess of NHS-Fluorescein (dissolved in DMSO; Thermo Fischer Scientific) and *in vitro* filamentation was analysed by epifluorescence microscopy. The NHS-Fluorescein dye was previously successfully used to visualize *in vitro* FtsZ and CCRP protein filaments (Camberg *et al.*, 2009; Springstein *et al.*, 2020b). And we note that the His_6_-tag did not impact the *in vitro* polymerization properties of the CCRP FilP (Javadi *et al.*, 2019), confirming the applicability of our approach.

## Supplementary Figures

**Supplementary Fig. 1:**
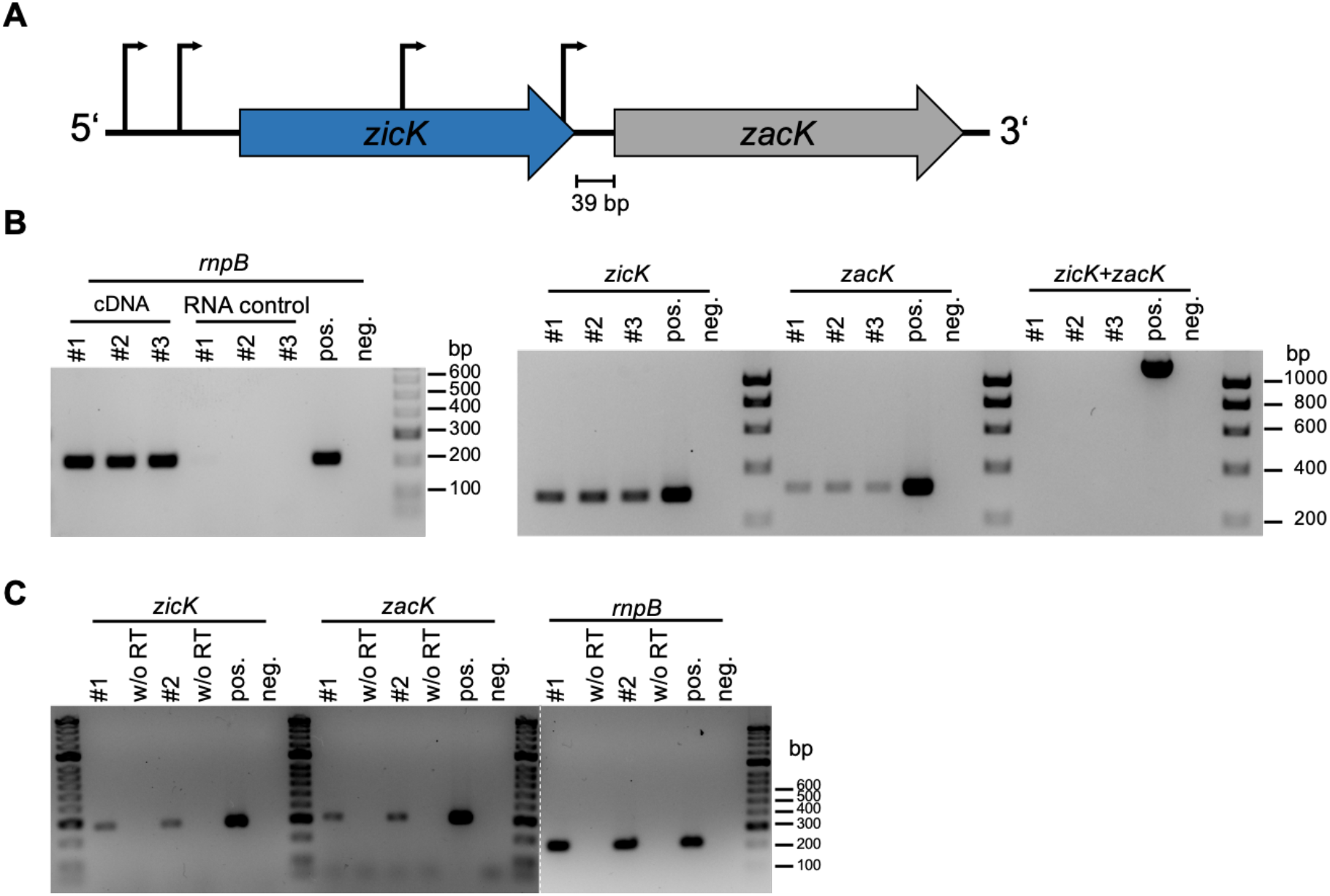
*zicK* and *zacK* are expressed at standard growth conditions. (**A**) Depiction of the genomic environment of *zicK* (blue) and *ZacK* (grey) within the *Anabaena* genome and their respective *in silico* predicted promoters depicted by black arrows (as predicted by BPROM (Solovyev and Salamov, 2011)). Promoters of *zicK* are predicted to reside 204 bp and 543 bp upstream of the open reading frame (ORF) and promoters of *zacK* are located 22 bp and 450 bp upstream of the ORF, thereby residing within the *zicK* ORF. (**B**,**C**) RT-PCR of whole cell RNA from *Anabaena* WT cultures grown in (**B**) BG11 or (**C**) BG11_0_ liquid medium from (**B**) three or (**C**) two independent biological replicates. Gene transcripts were verified using internal gene primers (Supplementary Table 4). As negative control (neg), PCR reactions were performed with water instead of cDNA or RNA and as a positive control (pos) *Anabaena* gDNA was included. 100 ng cDNA was used for each RT-PCR reaction. Absence of residual genomic DNA in DNase I-treated samples was verified with (**B**) 100 ng DNase I-treated RNA (RNA control) or (**C**) 100 ng DNase I-treated RNA that was subjected to cDNA synthesis reaction lacking reverse transcriptase (w/o RT).

**Supplementary Fig. 2:**
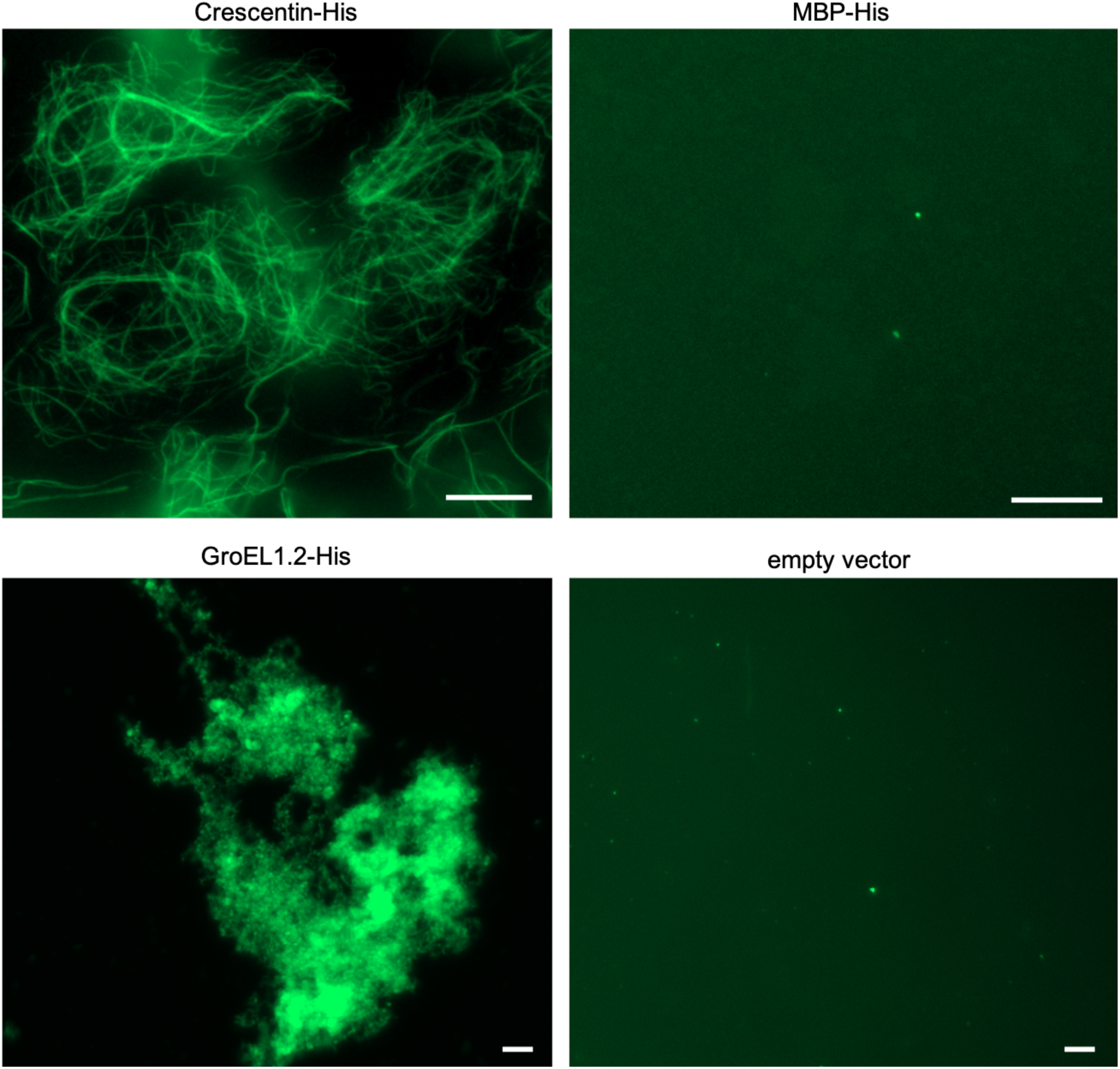
*In vitro* polymerization assay controls. NHS-fluorescein fluorescence micrographs of purified and renatured Crescentin-His, MBP-His and GroEL1.2 from *Chlorogloeopsis fritschii* PCC 6912 (0.5 mg ml^−1^ each) as well as purified cell-free extracts of *E. coli* BL21 (DE3) carrying empty vector (pET21a(+)) in HLB. While neither the cell-free extract containing empty vector nor the MBP protein formed any discernible structures *in vitro*, GroEL1.2 aggregates could be indicative for an uncontrolled oligomerization. We also observed similar clumps of protein aggregates from other *Anabaena* CCRPs that were negatively tested for *in vitro* polymerization. We therefore consider this *in vitro* behaviour a common property of putative oligomerizing proteins. Proteins and cell-free extracts (empty vector) were dialyzed in a stepwise urea-decreasing manner and stained with an excess of NHS-Fluorescein. Scale bars: 10 μm.

**Supplementary Fig. 3:**
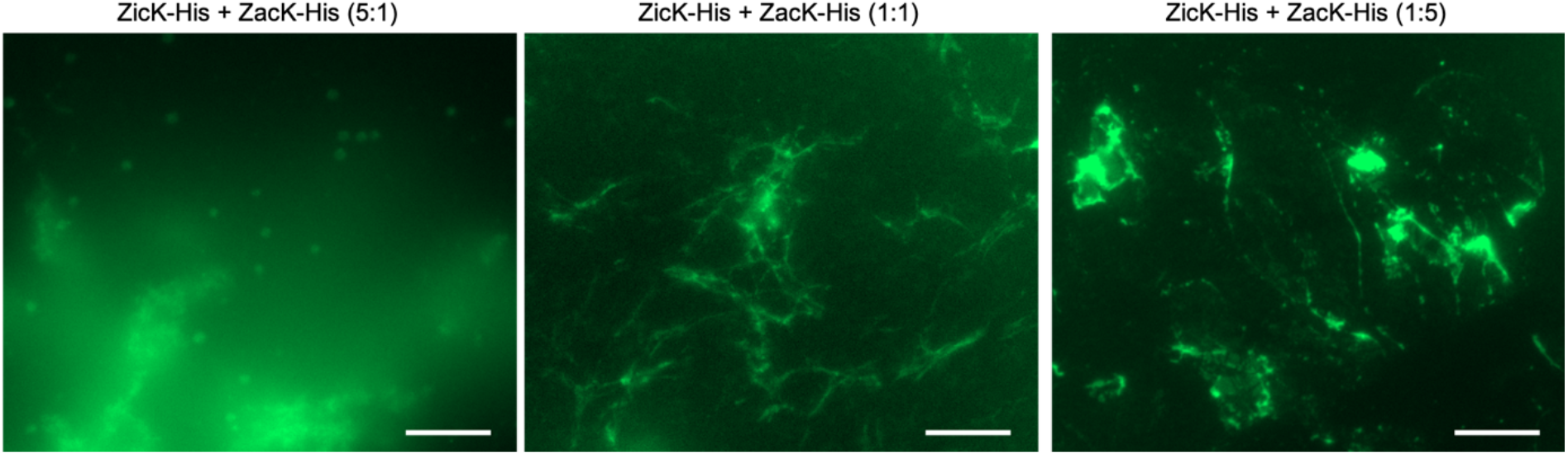
Co-polymerization of ZicK and ZacK is dosage-dependent. NHS-fluorescein micrographs of purified and co-renatured ZicK-His and ZacK-His in HLB renaturation buffer. ZicK-His and ZacK-His were combined in different ratios, either with a fivefold excess of ZicK-His (left image; corresponding to 0.25 mg ml^−1^ ZicK-His and 0.05 mg ml^−1^ ZacK-His), a fivefold excess of ZicK-His (right image; corresponding to 0.25 mg ml^−1^ ZacK-His and 0.05 mg ml^−1^ ZicK-His) or an equal concentration of ZicK-His and ZacK-His (central image; 0.25 mg ml^−1^ each). Proteins were dialyzed in a step-wise urea-decreasing manner and stained with an excess of NHS-Fluorescein. Fine heteropolymers only form when equal concentrations of ZicK-His and ZacK-His are present. In concert with the partial self-polymerization capacity of ZacK-His (Fig. 3A), certain filamentous structures are also detected in the ZacK-His excess samples. However, most protein still precipitated under those conditions. Scale bars: 10 μm.

**Supplementary Fig. 4:**
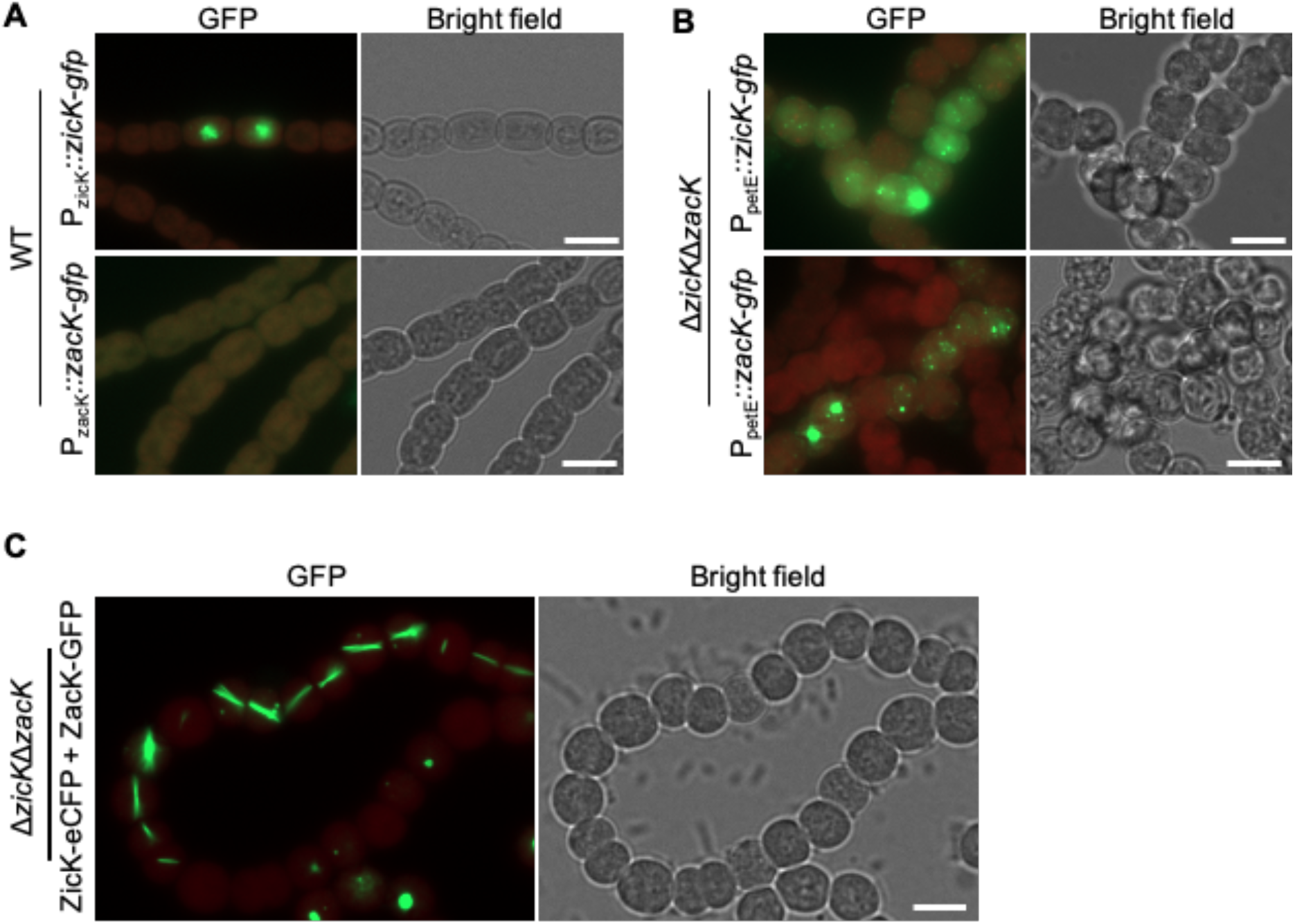
Heterologous expression of ZicK and ZacK. (**A**) Merged GFP fluorescence and chlorophyll autofluorescence (red) and bright field micrographs of *Anabaena* WT cells expressing ZicK-GFP or ZacK-GFP from P_zicK_ and P_zacK_. No expression of ZacK-GFP is detectable from P_zacK_ while expression of ZicK-GFP from P_zicK_ leads to similar patchy clumps within the cells as observed from P_petE_ in Fig. 4A. Note, we generally observed that the P_petE_-driven gene expression does not always lead to expression of the fusion protein in every cell under standard growth conditions. (**B,C**) Merged GFP fluorescence and chlorophyll autofluorescence and bright field micrographs of Δ*zicK*Δ*zacK* mutant expressing (**B**) ZicK-GFP or ZacK-GFP or (**C**) co-expressing ZicK-eCFP and ZacK-GFP from P_petE_. For expression of ZicK-GFP alone, BG11 plates were supplemented with 1 μM CuSO_4_. (**A**-**C**) These experiments show that expression of ZicK-GFP and ZacK-GFP or co-expression of ZicK-eCFP together with ZacK-GFP (Fig. 3A) from P_petE_ and their localization in *Anabaena* WT is not affected by ZicK or ZacK natively present in the WT background. Scale bars: (A-C) 5 μm.

**Supplementary Fig. 5:**
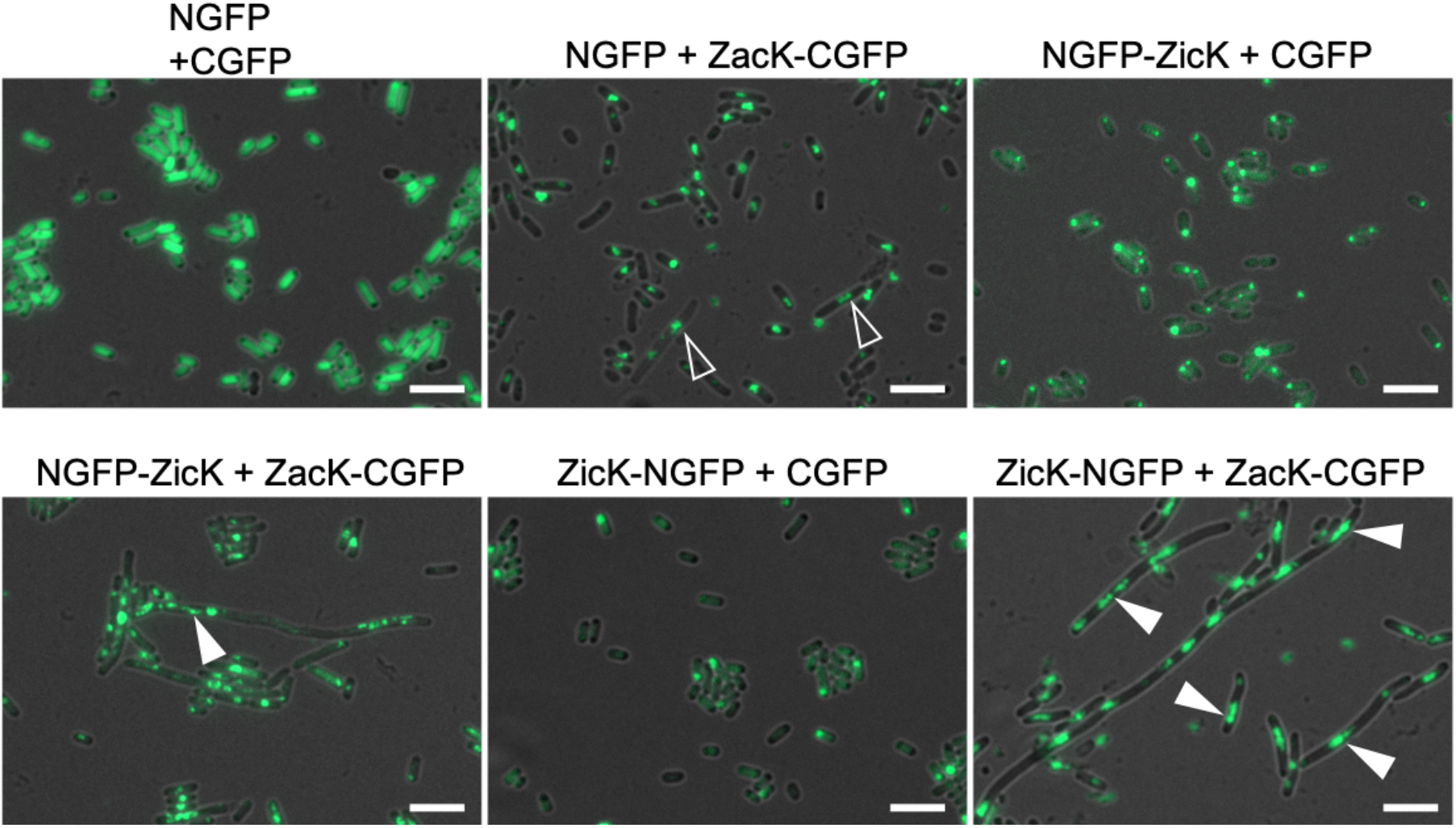
Host-independent heteropolymerization of ZicK and ZacK upon heterologous expression in *E. coli*. GFP-fragment reassembly assay. Merged GFP fluorescence and bright field micrographs of *E. coli* cells co-expressing NGFP (empty pET11a-link-NGFP) and CGFP (empty pMRBAD-link-CGFP), NGFP and ZacK-CGFP, NGFP-ZicK and CGFP, NGFP-ZicK and ZacK-CGFP, ZicK-NGFP and CGFP or ZicK-NGFP and ZacK-CGFP. Transparent triangles point to structures resembling ZacK-His *in vitro* polymers. White triangles indicate FilP-GFP-like (Bagchi *et al.*, 2008) filament-like structures that resemble structures indicated with translucent triangles but span longer distances. Co-expression of both, ZicK and ZacK leads to an elongated cell phenotype. FilP-like structures and elongated cells can already be seen upon co-expression of NGFP-ZicK with ZacK-CGFP but only the co-expression of ZicK and ZacK with C-terminal GFP-fragments leads to a clear filamentous cell phenotype and abundant intracellular filament-like structures. This suggests that the N-terminus of ZicK and ZacK is important for heteropolymerization. Note: we observed that the GFP-fragment reassembly assay is not suitable for the detection of protein-protein interaction strengths as even empty vector controls reconstitute the GFP protein, nonetheless, we employed it as a mean to localize the effect of co-expression of ZicK and ZacK in an entirely unrelated organism. Scale bars: 5 μm.

**Supplementary Fig. 6:**
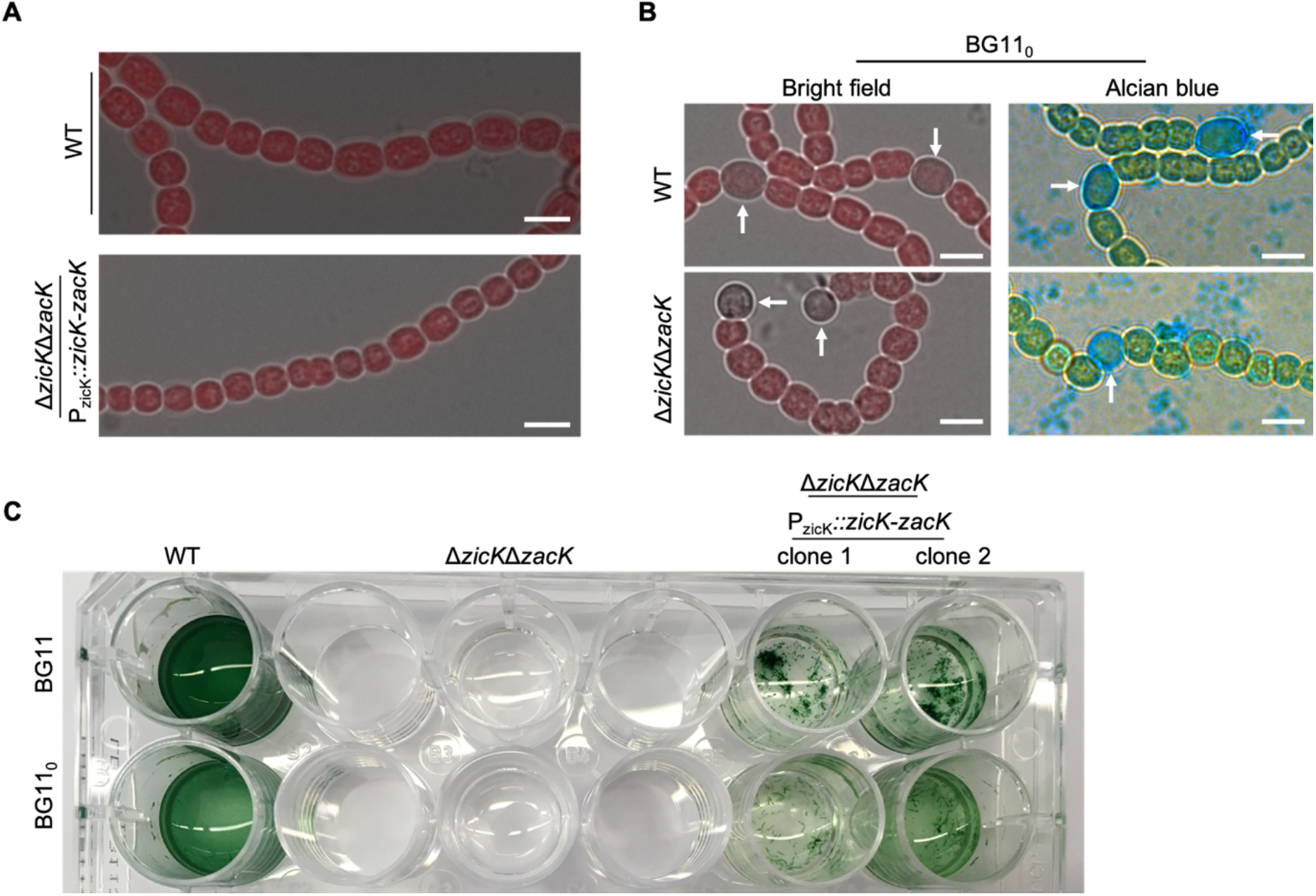
Mutant phenotype complementation and identification of heterocysts. (**A**) Morphological complementation of the ΔzicKΔ*zacK* mutant strain as a result of native expression of *zicK-zacK* from pRL25C. The capacity to complement the mutant phenotypes using the pRL25C plasmid shows that pDU1-based plasmids can be successfully employed to rescue WT phenotypes despite their variation in the relative copy number (Yang *et al.*, 2013). (**B**) Colour images of *Anabaena* WT and ΔzicKΔ*zacK* mutant strain grown on BG11_0_ plates and stained with alcian blue. (**A**,**B**) Scale bars: 5 μm. (**C**) Rescue of liquid growth of the Δ*zicK*ΔzacK mutant strains by expressing *zicK*-*zacK* from P_zicK_ from the replicative pRL25C plasmid.

**Supplementary Fig. 7:**
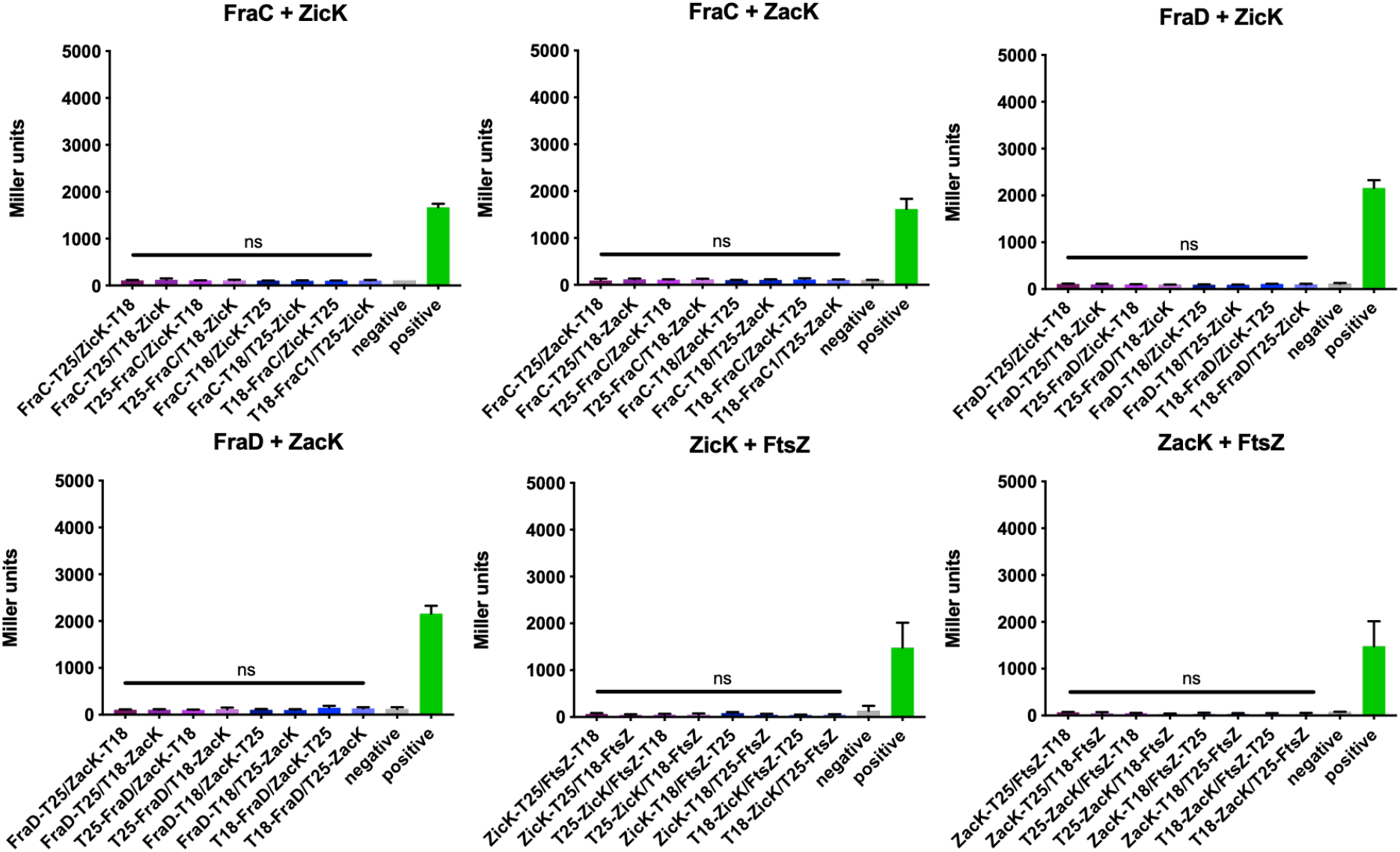
ZicK and ZacK do not interact with other components of the septal junctions besides SepJ. Beta-galactosidase assays of *E. coli* cells co-expressing indicated T25 and T18 translational fusions of all possible pair-wise combinations. Quantitative values are given in Miller units, and the mean results from three independent colonies are presented. Negative: N-terminal T25 fusion construct of the respective protein co-transformed with empty pUT18C. Positive: Zip/Zip control. Error bars indicate standard deviations (n = 3). *: P<0.05, **: P<0.01, ***: P<0.001, ****: P<0.0001 (Dunnett’s multiple comparison test and one-way ANOVA).

**Supplementary Fig. 8:**
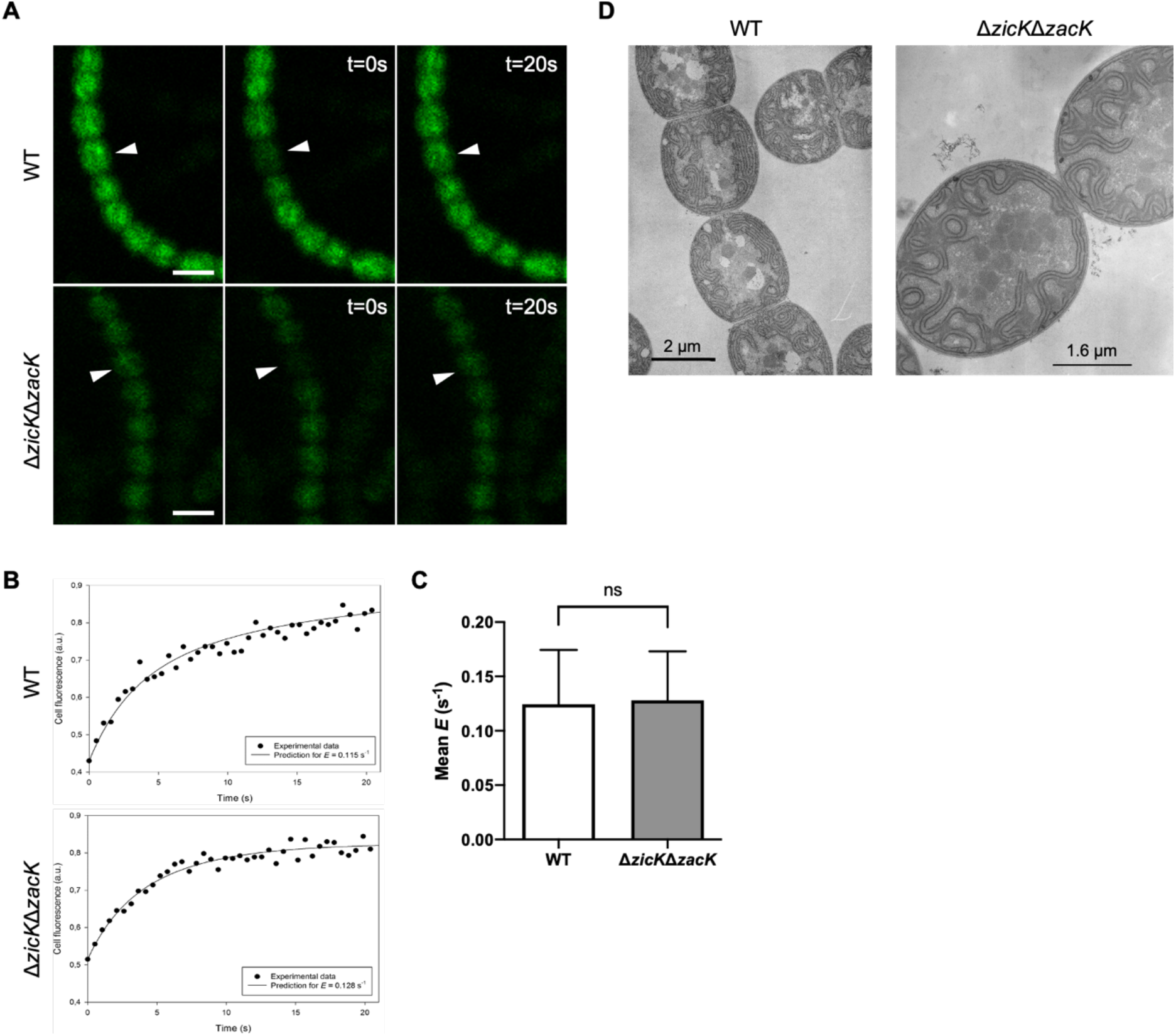
The Δ*zicK* Δ*zacK* mutant is not defective in intercellular transport and cellular ultrastructures. (**A**) Representative calcein fluorescence micrographs of FRAP experiments from calcein-labelled *Anabaena* WT and Δ*zicK*Δ*zacK* mutant grown on BG11 plates and (**B**) respective representative cell fluorescence recovery graphs. White triangles indicate bleached cells. Fluorescence images show respective cells prior bleaching, immediately after bleaching (t=0) and 20 seconds after bleaching (t=20s). Scale bars: 5 μm. **(C)** Mean exchange coefficients (E) of FRAP experiments from (**A**). Data present the number of recordings of bleached cells (*Anabaena* WT: n=21; Δ*zicK*Δ*zacK*: n=17). Values indicated with “ns” are not significantly different from the WT (using Student’s t-test). (**D**) Ultrathin sections of *Anabaena* WT and Δ*zicK*Δ*zacK* mutant strain grown on BG11 plates.

**Supplementary Fig. 9:**
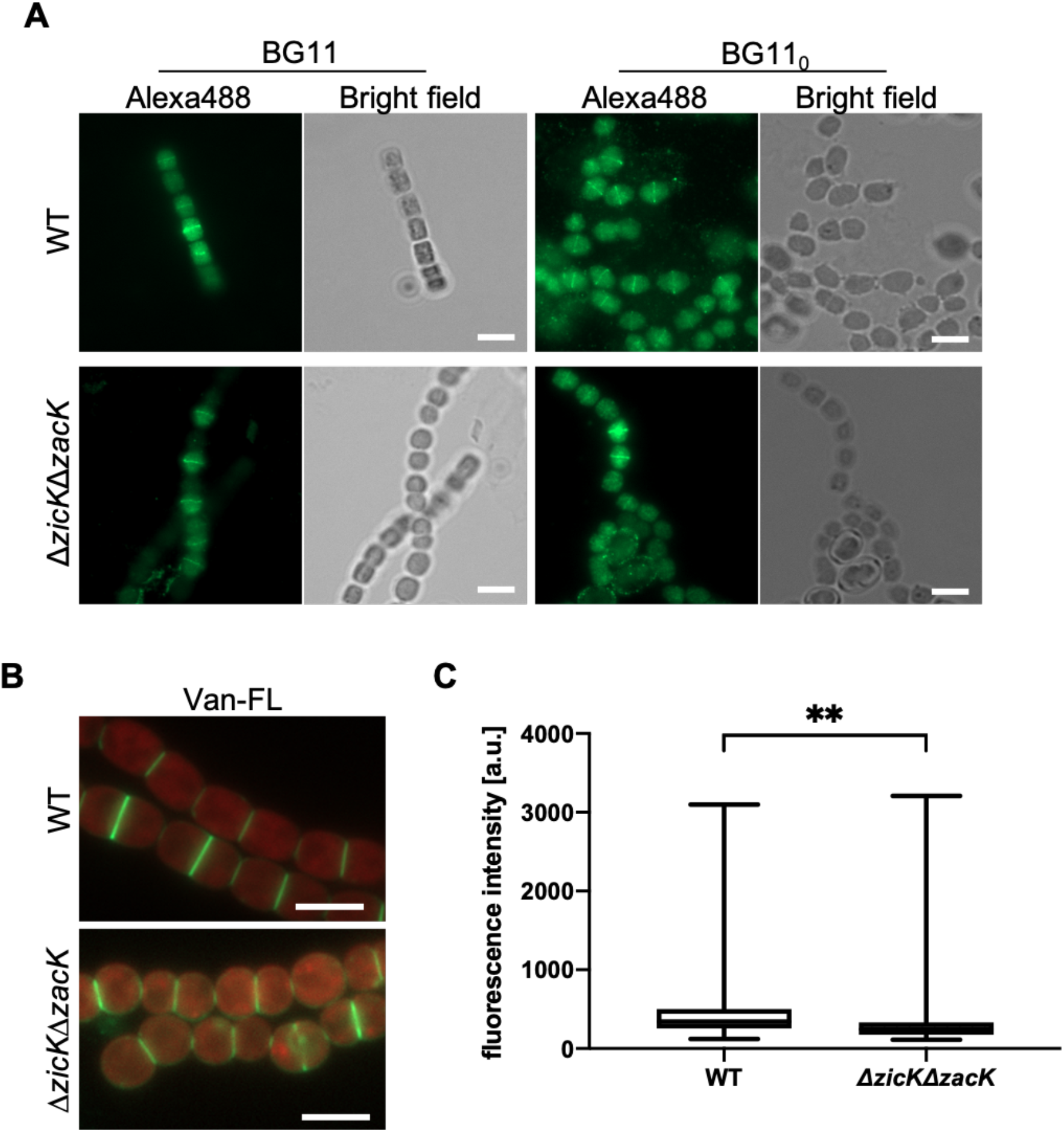
PG biogenesis and Z-ring placement are largely unaffected in the Δ*zicK* Δ*zacK* mutant. (**A**) Alexa Fluor-488 and bright field micrographs of Anabaena WT and Δ*zicK*Δ*zacK* mutant subjected to anti-FtsZ immunofluorescence. (**B**) Merged BODIPY™ FL Vancomycin (Van-FL) fluorescence and chlorophyll autofluorescence micrographs of *Anabaena* WT and the Δ*zicK*Δ*zacK* mutant stained with Van-FL. As a result of the low Van-FL staining and for better visibility, Van-FL fluorescence signal in Δ*zicK*Δ*zacK* mutant was artificially increased about twofold after image acquisition (note: this increase was not used for the fluorescence intensity measurement in (**C**)). Scale bars: (**A**,**B**) 5 μm. (**C**) Arithmetic mean fluorescence intensities of n=200 cell septa from (**B**). Values indicated with * are significantly different from the WT. **: P <.001, (Student’s t-test).

---

**Supplementary Table 1:**
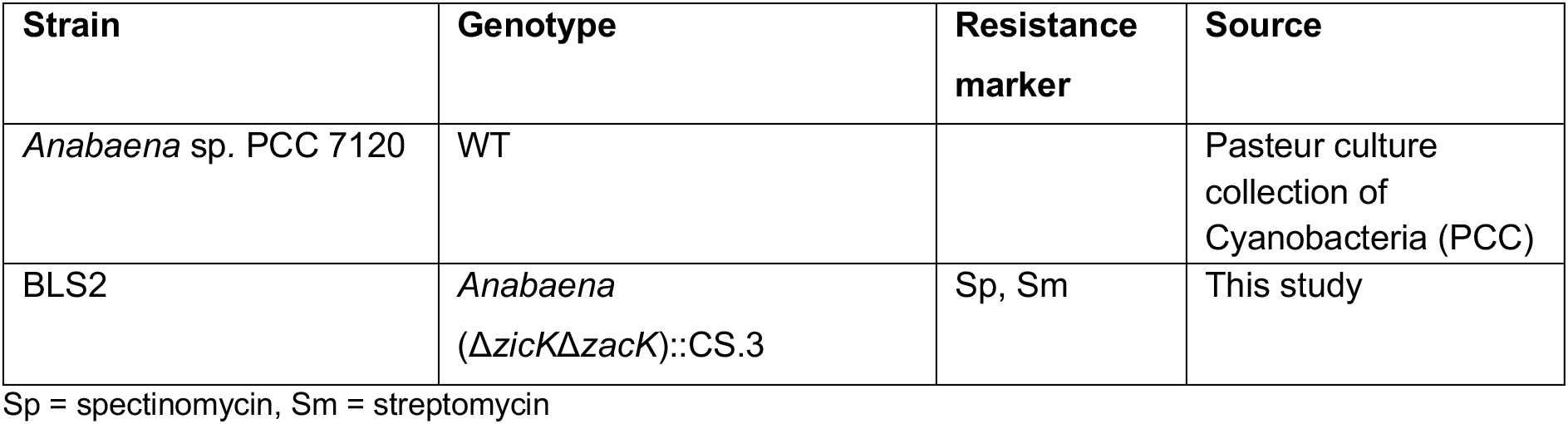
Cyanobacterial strains.

**Supplementary Table 2:**
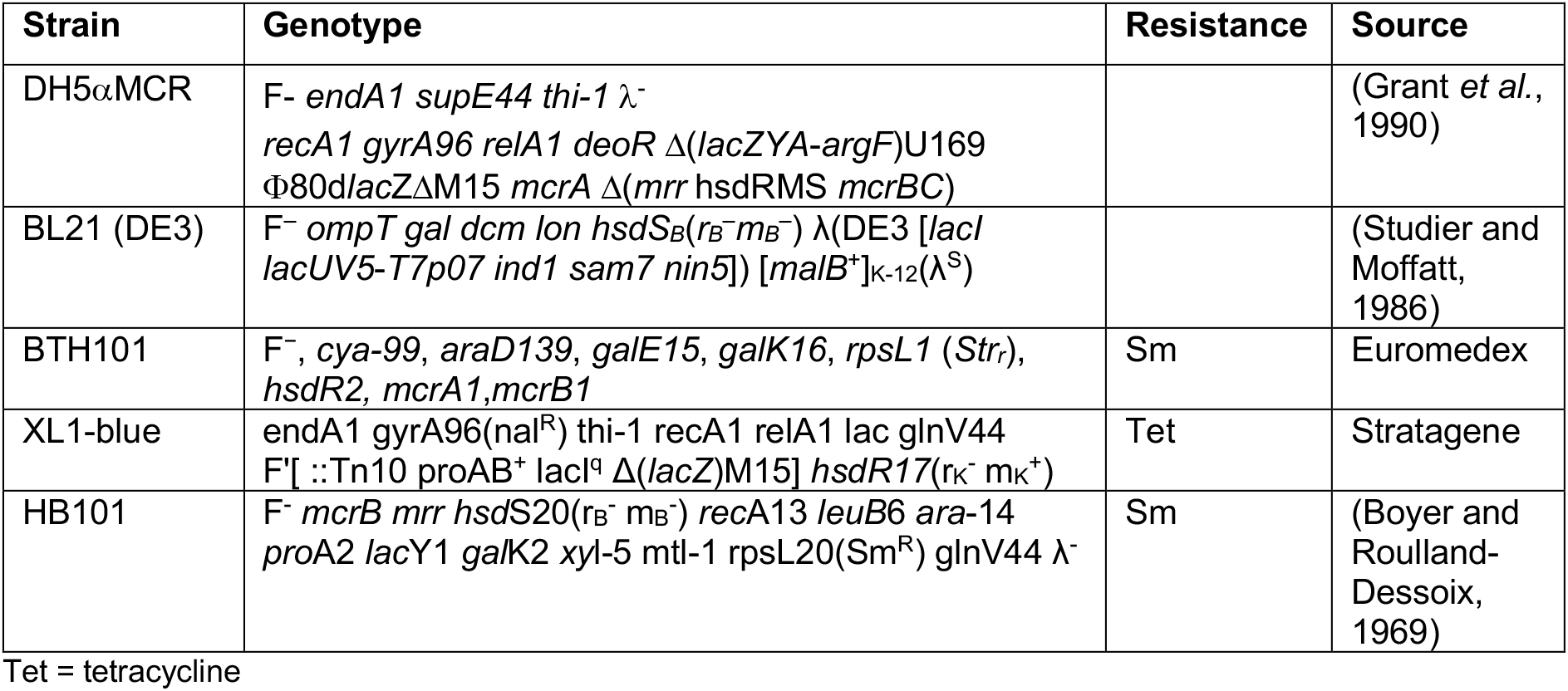
*E. coli* strains.

**Supplementary Table 3:**
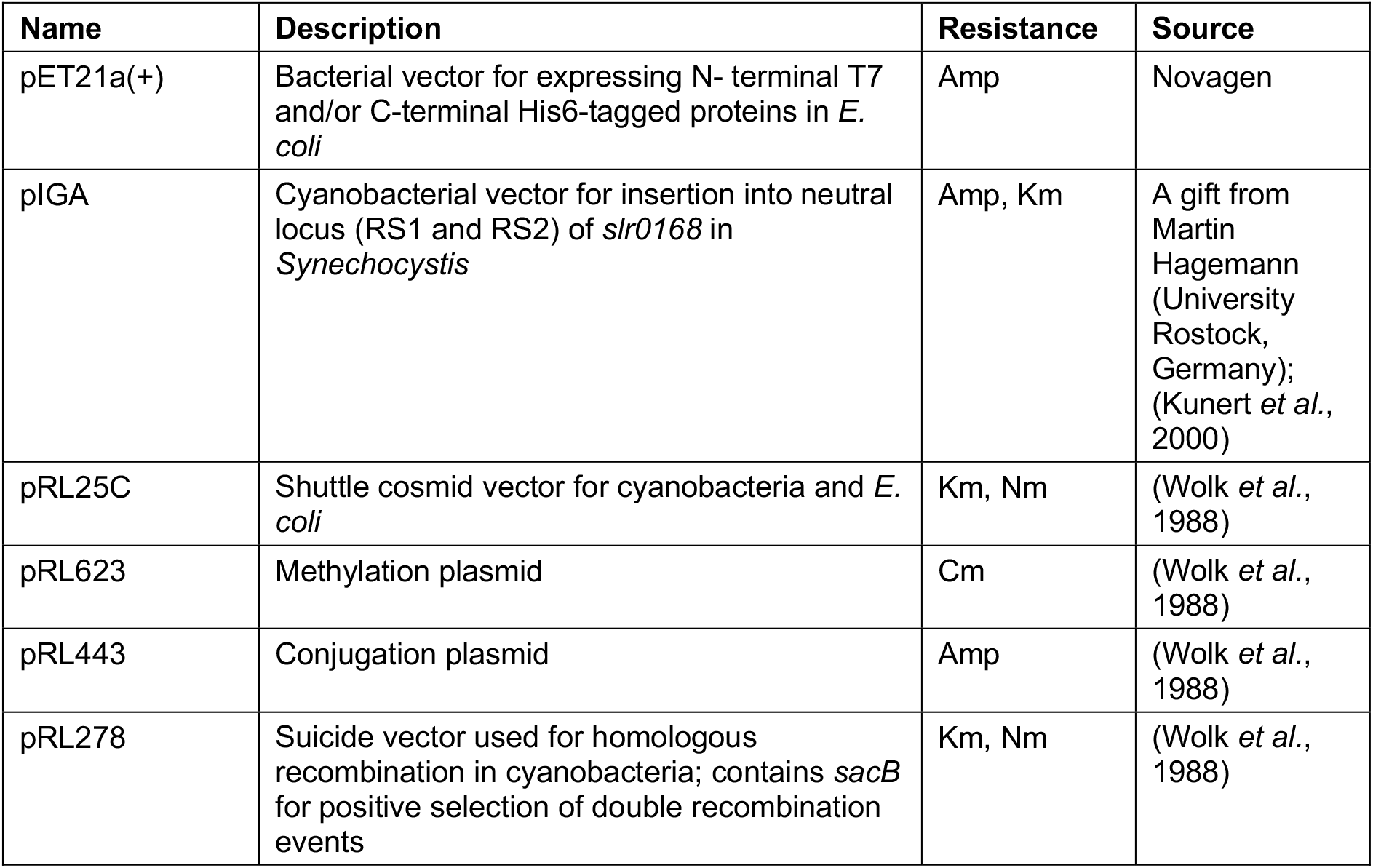
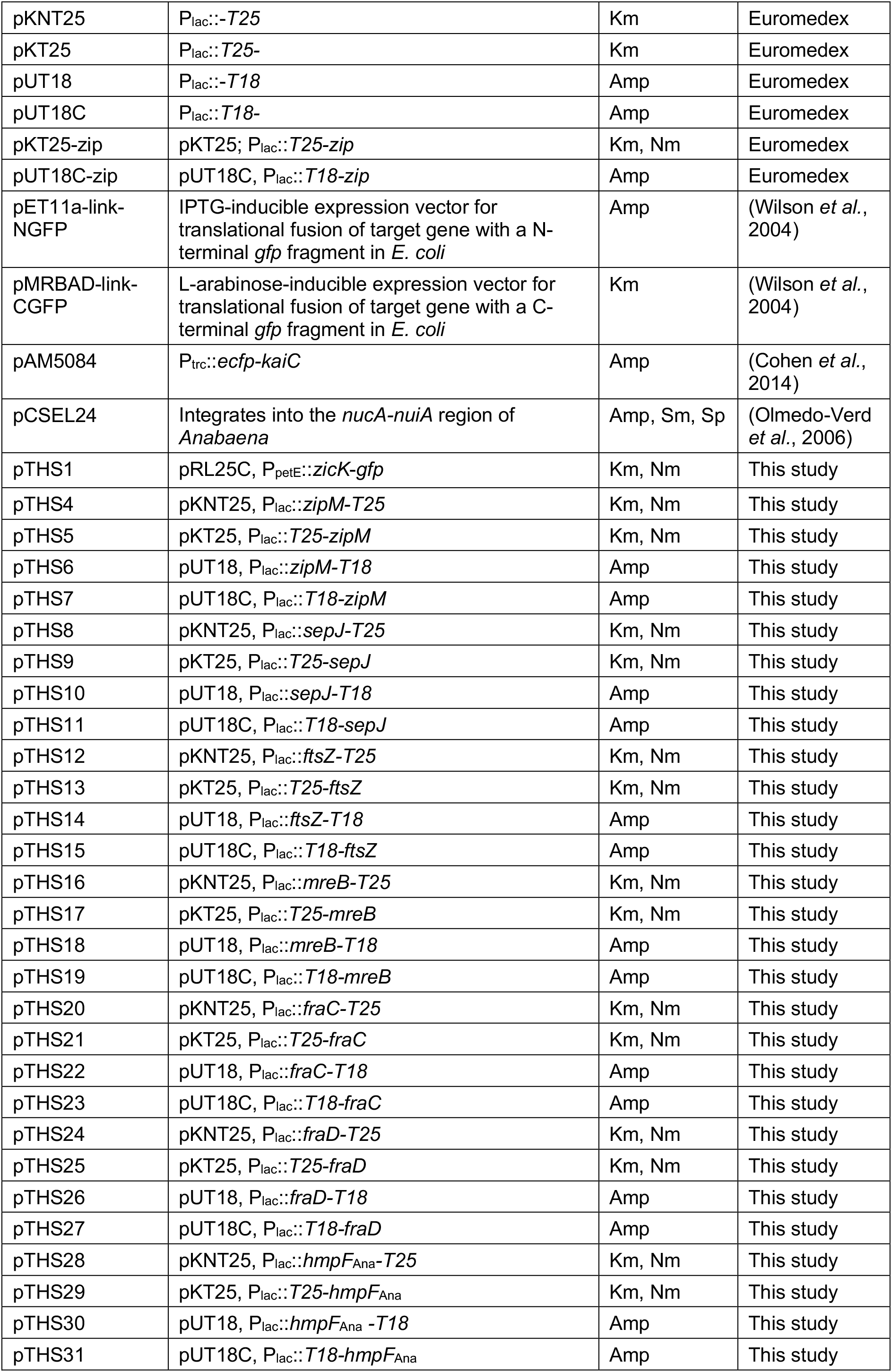
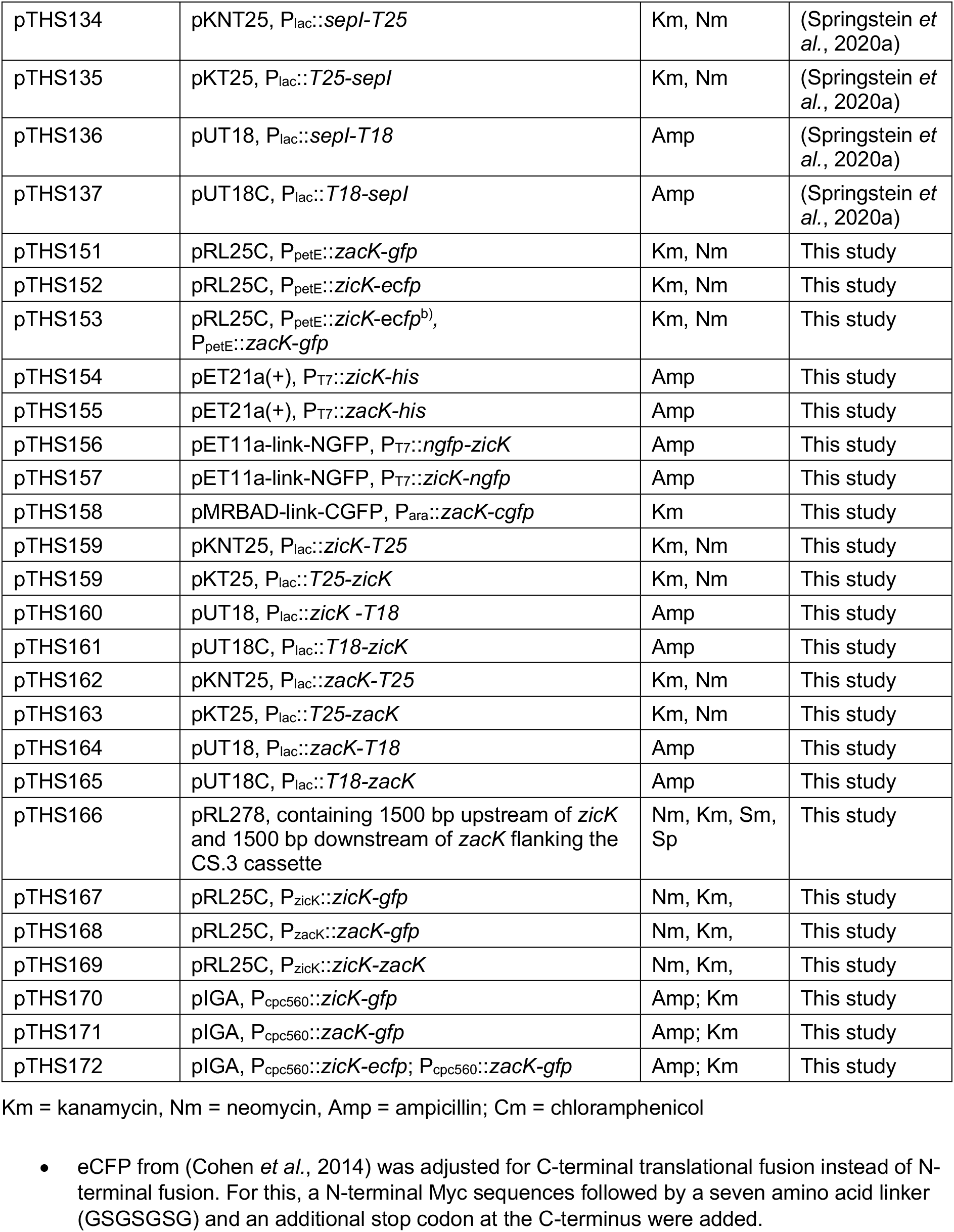
Plasmids.

**Supplementary Table 4:**
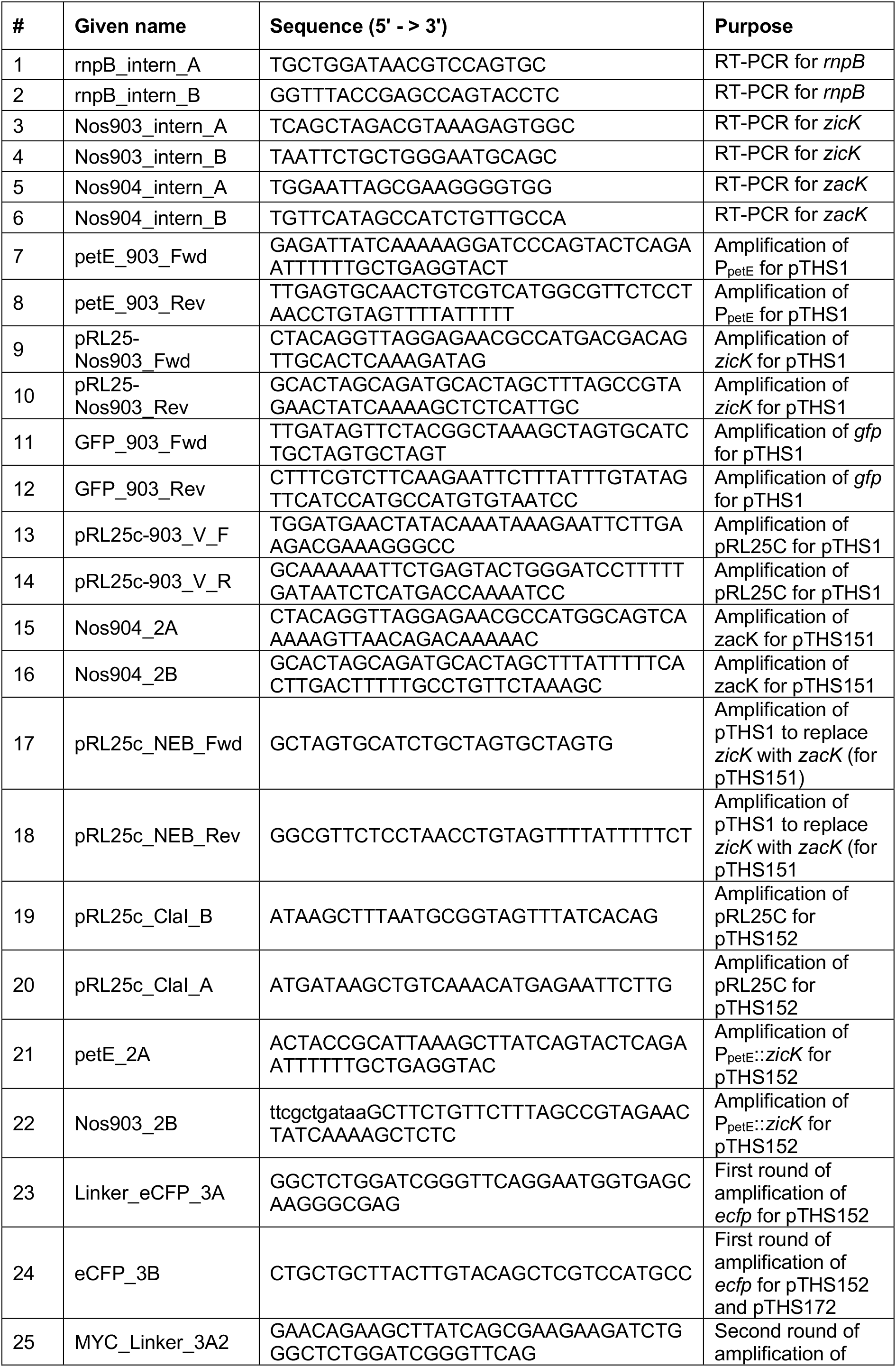
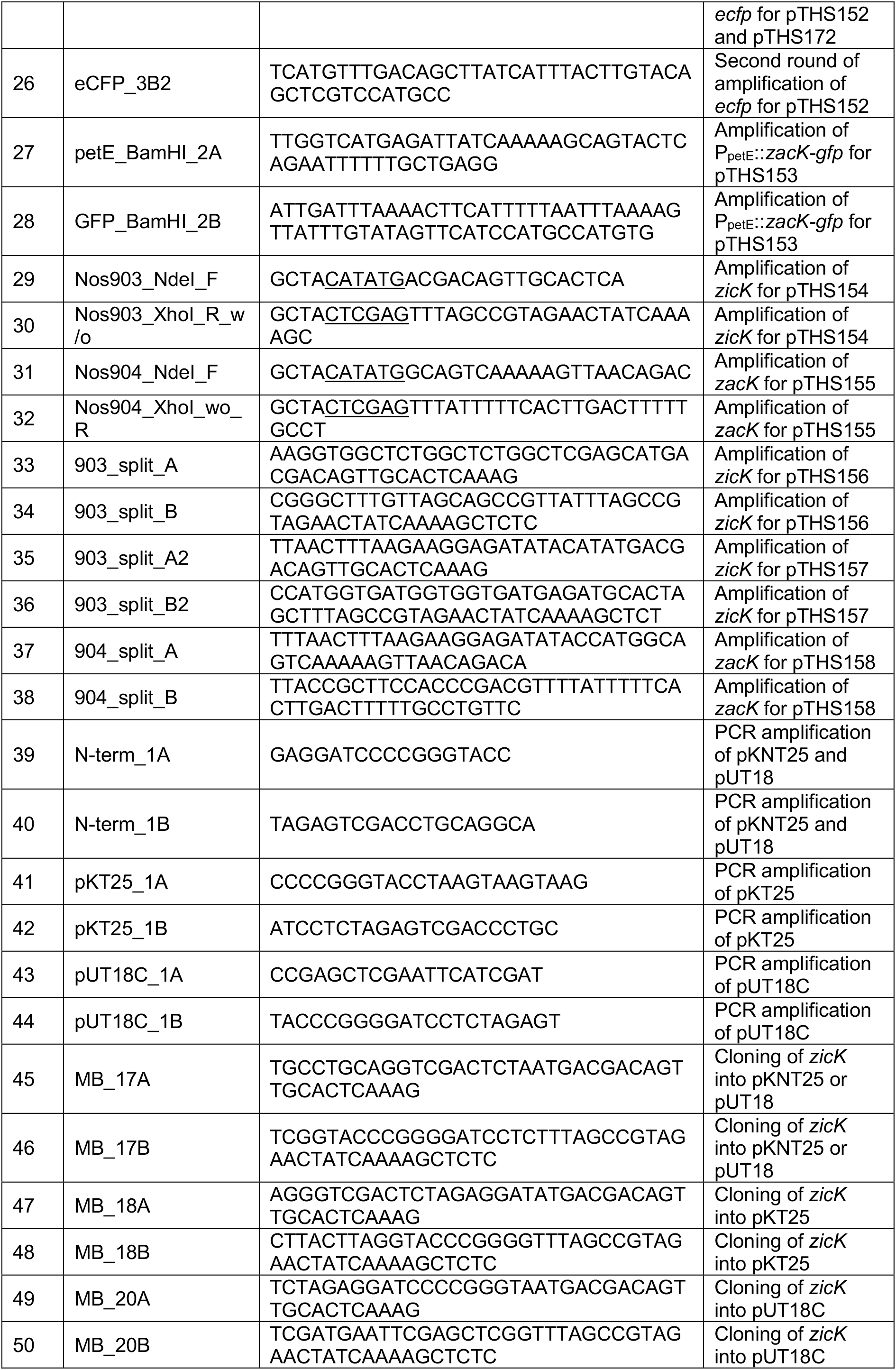
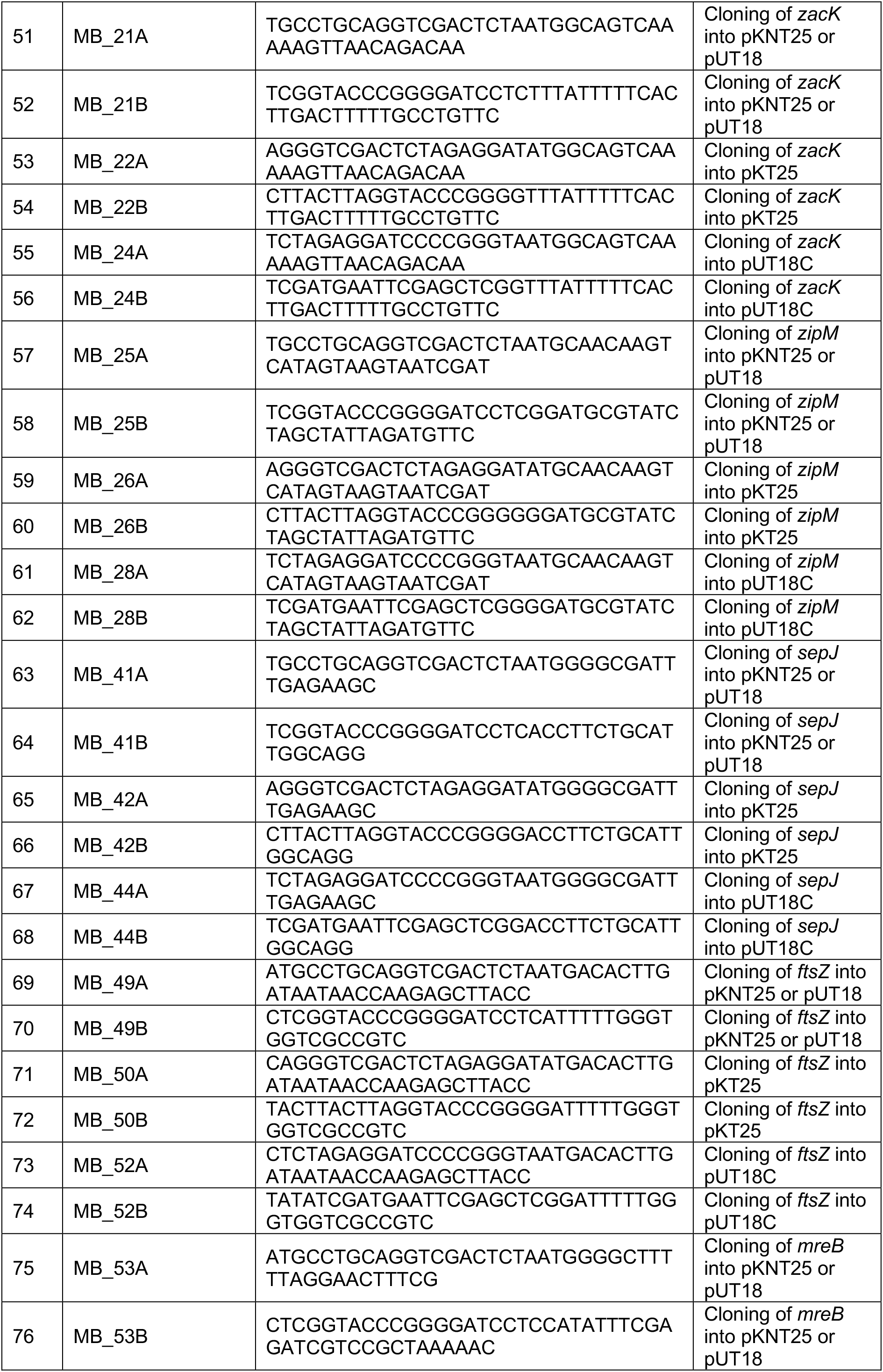
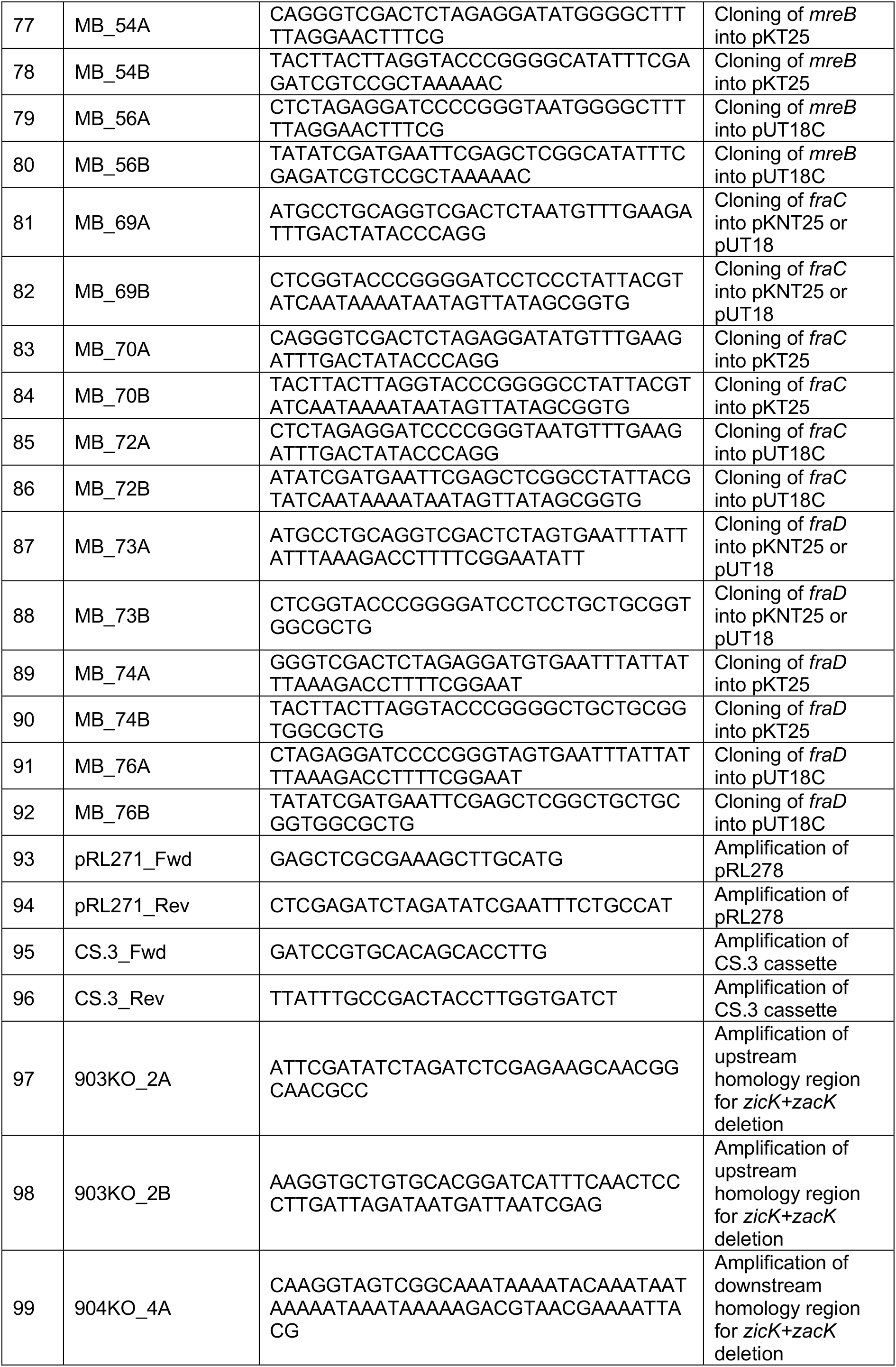
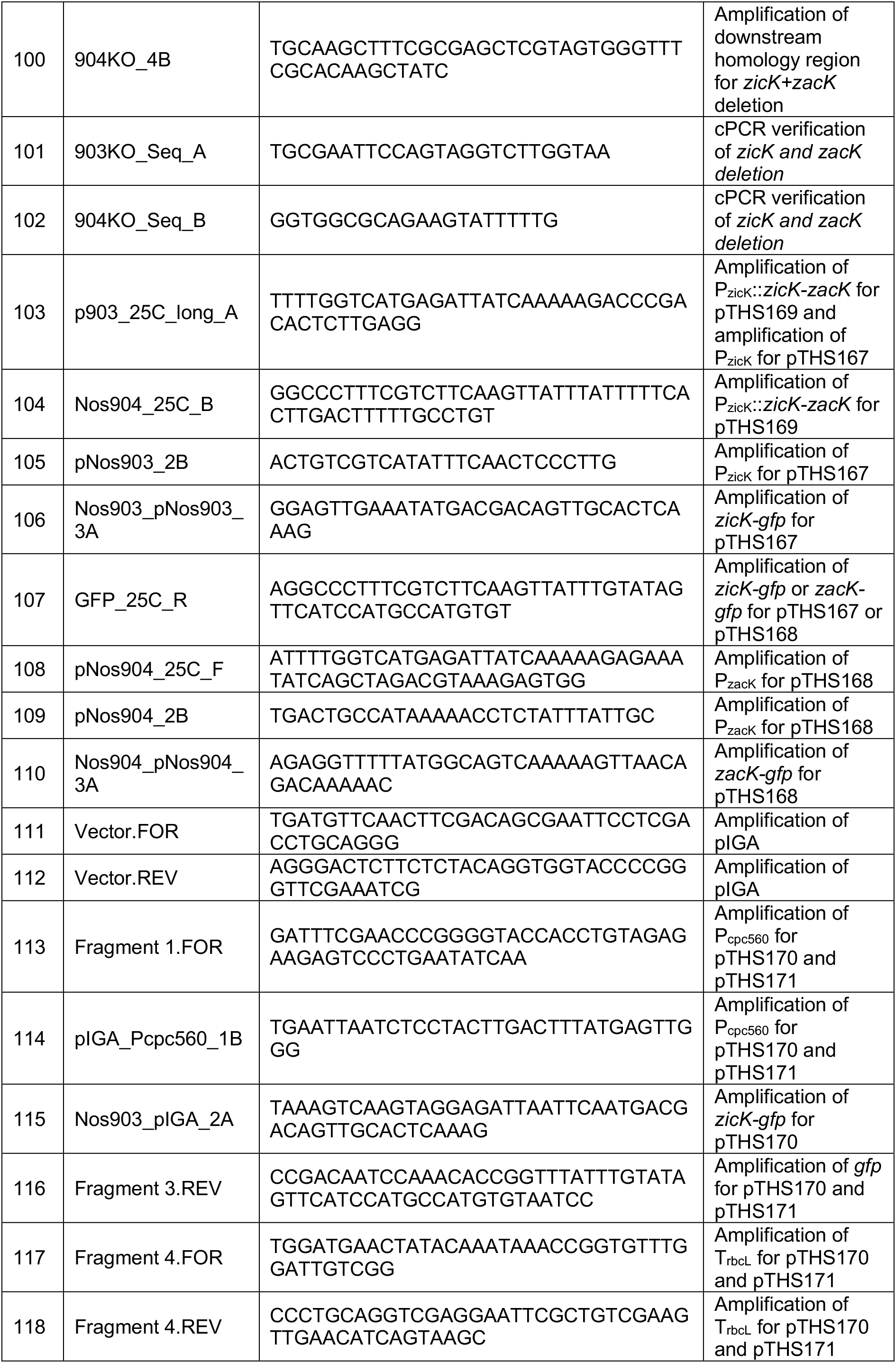
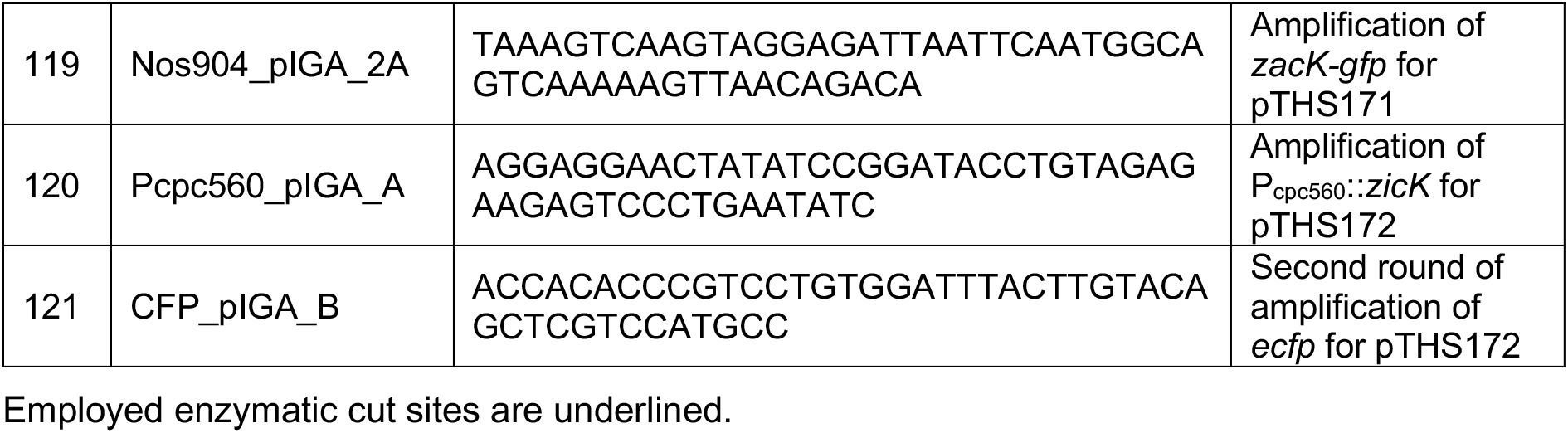
Oligonucleotides.

